# Precision-weighting of cortical unsigned prediction error signals benefits learning, is mediated by dopamine, and is impaired in psychosis

**DOI:** 10.1101/558478

**Authors:** J. Haarsma, P.C. Fletcher, J.D. Griffin, H.J. Taverne, H. Ziauddeen, T.J. Spencer, C. Miller, T. Katthagen, I. Goodyer, K.M.J. Diederen, G.K Murray

## Abstract

Recent theories of cortical function construe the brain as performing hierarchical Bayesian inference. According to these theories, the precision of cortical unsigned prediction error (i.e., surprise) signals plays a key role in learning and decision-making, to be controlled by dopamine, and to contribute to the pathogenesis of psychosis. To test these hypotheses, we studied learning with variable outcome-precision in healthy individuals after dopaminergic modulation with placebo, a dopamine receptor agonist bromocriptine or a dopamine receptor antagonist sulpiride (dopamine study n=59), and in patients with early psychosis (psychosis study n=74: 20 participants with First Episode Psychosis, 30 healthy controls and 24 participants with At Risk Mental State attenuated psychotic symptoms). Behavioural computational modelling indicated that precision-weighting of unsigned prediction errors benefits learning in health, and is impaired in psychosis. FMRI revealed coding of unsigned prediction errors relative to their precision in superior frontal cortex (replicated across studies, combined n=133), which was perturbed by dopaminergic modulation, impaired in psychosis, and associated with task performance and schizotypy (schizotypy correlation in 86 healthy volunteers). We conclude that healthy people, but not patients with first episode psychosis, take into account the precision of the environment when updating beliefs. Precision-weighting of cortical prediction error signals is a key mechanism through which dopamine modulates inference and contributes to the pathogenesis of psychosis.

## Introduction

A common theme in contemporary theories of brain function, ranging from perception (Rao & Ballard, 1999) to reinforcement learning (Mathys et al., 2011), is an emphasis on the critical role in inference played by predictions based on prior knowledge (Rao & Ballard, 1999; Friston, 2009; Bastos et al., 2012; Adams et al., 2013; Mathys et al., 2011; Clark, 2013 & 2015; Hohwy, 2013). According to these theories, predictions and incoming sensory input each have an associated precision (inverse variance), reflecting their confidence or reliability. Predictions and sensory input are thought to be compared against one other, generating a discrepancy signal termed the prediction error which indicates the difference between the expectation and sensory input. Such prediction error signals update prior beliefs in a manner that is weighted by their associated precision, such that more is learned from precise and reliable prediction errors compared to noisy and unreliable prediction errors (Friston, 2009; Bastos et al., 2012; Mathys et al., 2011). Several theorists have suggested that neuromodulator systems, including dopamine, play an important role in mediating the precision of these prediction errors, and that impaired precision-weighting of prediction errors (through dopaminergic or other neuromodulator dysfunction) may be part of the cascade that result in psychotic symptoms (Bastos et al., 2012; Adams et al., 2013; Friston, 2009; Fletcher & Frith 2009). However, to the best of our knowledge no direct evidence for these hypotheses exists. Specifically, whilst several neuroimaging studies have indicated abnormal brain prediction error signals in schizophrenia and related psychoses (Corlett et al 2007, Murray et al 2008, Schlagenhauf et al 2014, Ermakova et al 2018), none of these studies have addressed precision-weighting of prediction errors in patients.

In the context of reinforcement learning models, a distinction can be made between two types of prediction errors. First, the signed prediction error indicates whether an outcome is better or worse than expected, and thereby plays a crucial role in changing the value allocated to cues, thereby guiding future decisions (Schultz et al., 1997; O’Doherty et al., 2003 & 2004; Pessiglione et al., 2006; D’Ardenne et al., 2008; Diederen et al., 2016 & 2017, Tian et al., 2016). A second type of prediction error, the unsigned prediction error, signals the degree of surprise, without indicating valence (better/ worse than expected). In addition to signed prediction errors, unsigned prediction error are included in various reinforcement learning models to control how much should be learned from new information. Large unsigned prediction errors signal that the brain’s model of the world is inaccurate, thereby increasing the amount that is learned from new information. This can be achieved in various ways, including a non-Bayesian approach by using a dynamic learning rate parameter (Pearce & Hall, 1982; Sutton & Barto, 1998) or a Bayesian approach by decreasing the precision of prior beliefs (Courville et al., 2006; Gershman, 2015) across different levels in the hierarchy so that new sensory information has more of an impact on learning (Mathys et al., 2011). In these hierarchical models both signed and unsigned prediction errors are weighted by their precision. Whilst evidence has been provided for a dopamine-mediated precision-weighted signed prediction error in learning (Diederen et al., 2015, 2016 & 2017), no such evidence exists for dopaminergic modulation of the precision-weighting of unsigned prediction errors. This is despite many computational theorists hypothesizing a role for neuromodulator systems in precision-weighting of unsigned prediction errors (Bastos et al., 2012; Adams et al., 2013), and that the dysfunctional precision-weighting is a key contributor to the pathogenesis of psychosis (Fletcher & Frith 2009; Adams et al 2013, Sterzer et al 2018, Heinz et al 2018).

Here we therefore studied whether unsigned prediction errors are indeed coded relative to their associated precision; whether dopamine modulates the precision of these prediction errors; where precision-weighted prediction errors are represented in the cortex and whether and how precision weighting of cortical prediction error signals is disrupted in psychosis. Our methods elicit reliable measures of prediction error, as we use a task where prediction error is directly observable, rather than inferred as a latent variable as is common in many paradigms. To probe the role of precision-weighted prediction errors in learning, and the influence of dopamine, we employed pharmacological modulation in healthy volunteers, combined with fMRI, associative learning and computational modelling. We next examined how individual differences in computational learning signals and brain precision weighting signals relate to clinical psychosis, and psychotic-like thinking in health (schizotypy).

## Methods

### Participants and intervention - dopamine study

59 healthy volunteers completed the pharmacological fMRI study (see Diederen et al., 2017 and Supplement); all provided written informed consent. Prior to scanning, participants received a single dose of the D2-antagonist sulpiride (600mg), the dopamine agonist Bromocriptine (2.5 mg), or placebo, in a double-blind fashion. The study received NHS research ethics approval.

### Participants - psychosis study

Healthy volunteers (HCS, n=30, average 22.6 years, 15 female) without a history of psychiatric illness or brain injury were recruited as control subjects. Healthy volunteers did not report any personal or family history of neurological, psychiatric or medical disorders. As in our previous work (Ermakova et al., 2018), we recruited participants with first episode psychosis with active delusions or hallucinations (PANSS P1 or P3 >2) (FEP, n=20 average 24.8 years, 6 female) or at-risk of psychosis (At Risk Mental States ARMS, n=24, average 21.5 years, 8 female) were recruited from the Cambridgeshire early intervention in psychosis service (Table 2). In addition, *potential* at-risk participants were identified on the basis of belonging to a help-seeking, low-mood, high schizotypy sub-group from the Neuroscience in Psychiatry Network (NSPN) cohort (Davis 2017) or through advertisement via posters displayed at the Cambridge University counselling services. Individuals at-risk for psychosis met At-Risk-Mental-State (ARMS) criteria on the CAARMS interview in the past six months (Yung et al. 2003). All participants gave written informed consent. The study received NHS research ethics approval.

**Table 1:** Demographics for dopamine study.

| Dopamine study |  |  |  |  |  |  |  |
| --- | --- | --- | --- | --- | --- | --- | --- |
|  | Placebo |  | Sulpiride (antagonist) |  | Bromocriptine (agonist) |  | p-value |
| N | 20 |  | 20 |  | 19 |  |  |
| Male | 9 (11) |  | 12 (8) |  | 10 (9) |  |  |
|  | Mean | SD | Mean | SD | Mean | SD |  |
| Age | 23.9 | 4.8 | 24.8 | 4.5 | 23.7 | 4.3 |  |
| Reverse digit span | 6.2 | 1.5 | 5.7 | 1.5 | 6.8 | 3.8 | *p*=.40 |
| Schizotypy | 16.6 | 10.1 | 15.4 | 11.5 | 15.6 | 11.2 | *p*=.99 |

**Table 2:** Demographics for psychosis study.

| Group | Healthy controls (HCS) |  | At risk mental state (ARMS) |  | FEP (First episode psychosis) |  |  |
| --- | --- | --- | --- | --- | --- | --- | --- |
| N | 30 |  | 24 |  | 20 |  |  |
|  | Mean | SD | Mean | SD | Mean | SD | p-value |
| Age | 22.6 | 3.5 | 21.0 | 3.5 | 24.9 | 5.2 | $p=.009$ |
| Male | 16 | | 18 | | 17 | | $p=.044$ |
| IQ | 118.8 | 10.5 | 118.7 | 11.7 | 106.3 | 20.2 | $p=.006$ |
| PANSS positive | 7.2 | 0.8 | 14.0 | 3.00 | 20.6 | 5.4 | $p<.001$ |
| PANSS negative | 7.2 | .9 | 13.5 | 6.2 | 15.5 | 8.4 | $p<.001$ |
| Taking antipsychotic medication | 0/30 | | 4/24 | | 12/20 | | $p<.001$ |
| Antipsychotic dosage | 0 | 0 | 55.5 | 64.3 | 353.5 | 184.54 | $p=.008$ |

### fMRI task design

The task (see Figure 1 and Supplement) consisted of three sessions of 10 minutes each. Rewards were drawn from six different pseudo-Gaussian distributions that differed with respect to their precision (i.e. inverse variance) and expected value (i.e. mean of the distribution; EV). For the dopamine study, the standard deviations from the distributions were 5, 10 & 15, corresponding to precision of .04, .01 and .004. For the psychosis study, the standard deviation of the distributions were either 5 or 15, corresponding to precision of .04 and .004. Distributions were counterbalanced to ensure that the two conditions within each session differed with respect to the mean of the distribution and precision. Conditions were presented in short blocks, each including 4-6 trials. Each distribution consisted of 31 trials, resulting in 62 trials per session. Participants were informed beforehand that each distribution, of which two per session (run), had a different level of precision, which could be one of three levels: low, medium or high precision, corresponding to precisions of 0.004, 0.01 and 0.04 although the exact precisions were not revealed to the participants. Furthermore, participants were instructed that the two means within a session would be different from each other. In the psychosis study there was no medium condition. A cue informed the participants from which of the distributions (high, medium or low) in that block the upcoming reward was being drawn; then participants were required to predict the magnitude of the upcoming reward and received feedback after a delay. Optimal performance on this task thus required the participant to estimate the mean of the distribution from which the rewards were drawn. For MRI acquisition details see Supplementary methods.

**Figure 1:**
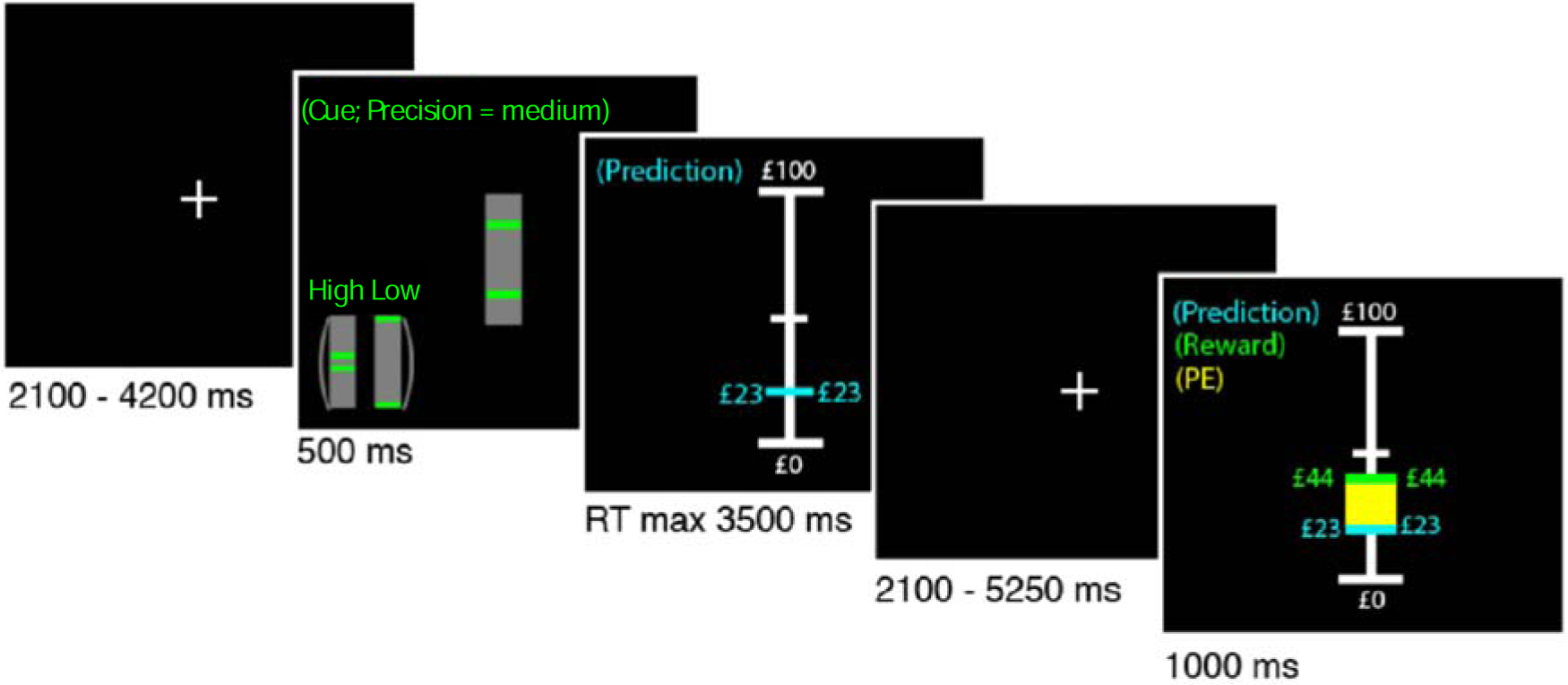
Example of a trial. The participants were instructed to learn the mean of a reward distribution. First a fixation cross was presented after which the participants were informed about the standard deviation (which indicated the precision) of the reward distribution. Subsequently the participants were asked to make a prediction regarding the upcoming reward, which was presented to the participant in combination with the prediction error (in yellow) after an anticipation period.

### Behavioural analysis and computational modelling

The mean performance error – the absolute value of (actual mean – predicted mean) – was our index of performance, which we compared across groups. We also fitted several reinforcement learning models to participants’ prediction sequences (see Supplementary methods). In brief, each model used a common updating rule in which predictions on a given trial depended on the prediction error and the learning rate on the previous trial. We implemented a Rescorla-Wagner (RW) reinforcement learning model with a fixed learning rate (Rescorla & Wagner 1972) and a Pearce-Hall (PH) model with a trial-wise, dynamic, learning rate, which prescribes higher weighting of prediction errors (i.e., more learning) at the start of a task session compared to later trials (Pearce & Hall 1982). In uncertain environments, it is more optimal to decrease the weighting of prediction errors as learning progresses (once participants become more certain of their predictions) as prediction errors will continue to occur as a result of the imposed uncertainty. We additionally explored whether scaling prediction error to the reliability of the environment (i.e., precision– weighting) benefitted learning by comparing models that scaled the prediction error term and models that did not. This results in 6 models: (1) simple RW, (2) RW with a scaled prediction error term, (3) Pearce-Hall model (decaying learning rate), (4) Pearce-Hall model with scaled prediction error term, (5) Pearce-Hall model with individual estimates of scaling term, (6) Pearce-Hall model with individual estimates of separate signed and unsigned prediction error scaling.

### Brain imaging analysis

We modelled the onsets of the cue and the outcome as events (i.e. delta functions of zero duration) and the onset of the prediction event (i.e., when participant could start making their prediction) as a single epoch lasting until they indicated their prediction. Each predictor was convolved by the standard canonical haemodynamic response function in SPM8. We used parametric modulation to identify neural correlates of unsigned prediction error responses by specifying for all outcome events the unsigned prediction errors. It is important to note that the prediction errors used in these analyses are simply the absolute difference between predicted reward and received reward. As such, the prediction errors did not dependent on the behavioural modelling, and could therefore not be influenced by any differences in the best-fitting model between groups. In a separate analysis, we also explored the coding of signed prediction errors in the psychosis study. The effect of dopaminergic drugs on the precision-weighting of signed prediction errors has been published before (Diederen et al., 2017). Reward events were separately modelled for the different precision conditions to test for differences in precision-weighting of prediction errors as evidences by different sizes of slopes for the coding of unsigned prediction errors under different levels of certainty. Contrasts were created on the 1^st^-level. As we were interested in the effect of precision but not mean reward, we collapsed all the different means for each precision condition, so there were two or three precision conditions in the psychosis study and dopamine study respectively, to be taken to the 2^nd^-level (i.e., group-level): see Supplement for details of group-level analysis.

## Results

### Study 1: Dopamine modulation study

#### Environmental precision and dopamine D2 receptor antagonism modulate task performance

Participants’ performance (the distance between the participants prediction and the mean of the distribution) improved when the precision of the reward distributions increased, and reduced average performance under sulpiride (Figure 2); Trial-by-trial analysis revealed lower performance in low-precision conditions at the start of the experiment, but no significant group differences (Figure 2).

**Figure 2:**
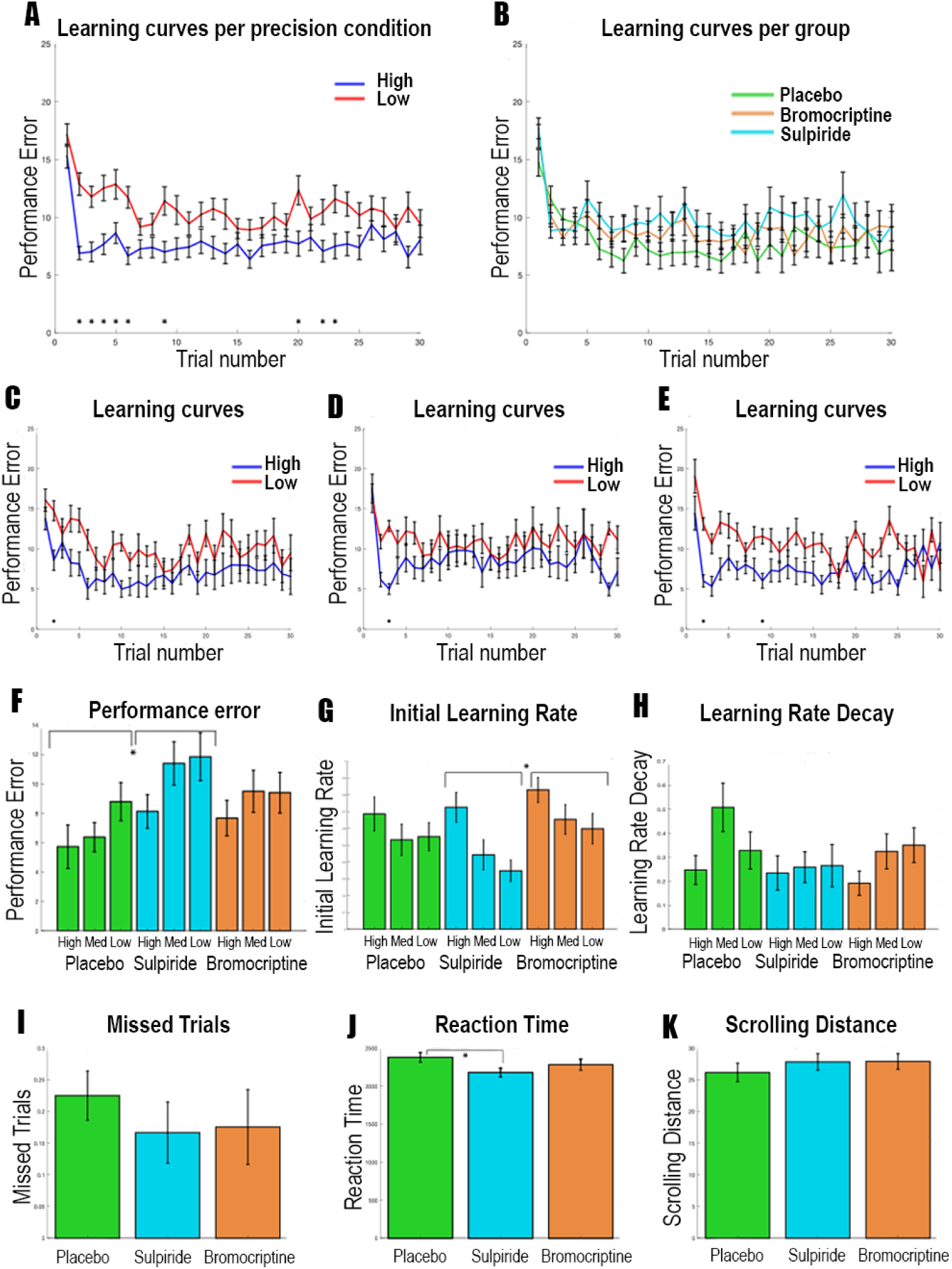
A. Behavioural results for the dopamine study. A-E display the average learning curves, reflecting the absolute distance between the actual mean of the distribution and the participants estimate of the mean distribution over 30 trials averaged over the 3 sessions for each participant. The distance between the prediction and the actual mean of the distribution defines performance error. Therefore, lower values of performance error are better. Asterisks indicate Bonferroni corrected significant differences across conditions. A: Performance was significantly better in the high precision condition compared to the low precision condition when combing all participants, especially at the beginning of the experiment. B: There were no clear differences between groups when analysing trial-by-trial performance. C-E: The placebo (C), Sulpiride (D) and Bromocriptine (E) group showed only a significant difference between the precision conditions one or two trials in the beginning of the experiment. F: Averaging performance error across all trials, we see overall better performance in the placebo condition compared to the Sulpiride condition, and higher performance in more precision conditions. G: Initial learning rates were higher in the Bromocriptine condition compared to the Sulpiride condition. H: Precision and group did not affect the learning rate decay parameter. I: No differences were found in number of missed trials, and K: scrolling distance. J: Reaction times were quicker in the Sulpiride group. Error bars represent standard error of the mean.

#### Reinforcement learning modelling of behavioural data indicates precision-weighted unsigned and signed prediction errors

Since formal learning models like the Pearce-Hall model suggest that unsigned prediction errors increase learning, we expect an interaction between unsigned and signed prediction error on participants’ trial to trial updates. The unsigned*signed prediction error interaction term was highly significant in predicting updates (F{1,10763}=51.7, p<.0001), demonstrating the importance of unsigned prediction errors in learning (Supplementary information). In all three medication groups a Pearce-Hall model with separately estimated precision weighted signed and unsigned prediction errors best predicted behaviour (Supplementary results). These results indicate that both unsigned and signed prediction errors are precision-weighted to facilitate efficient learning under uncertainty.

#### Unsigned prediction errors are coded in the Superior Frontal Cortex (SFC) and pre SMA/dACC

We next explored where unsigned prediction errors are coded in the brain, in order to find the region of interest for our further analyses focussing on the effect of dopamine on the precision-weighting of prediction errors. Whilst correcting for whole brain comparisons, unsigned prediction errors were coded in the frontal, parietal and occipital cortices (Supplementary Table 2, Figure 3). We used the left and right SFC and dACC clusters as ROIs to take forward our analysis of dopaminergic effects on precision weighting. Secondary analyses examined these effects in occipital and parietal regions (Supplementary materials).

**Figure 3:**
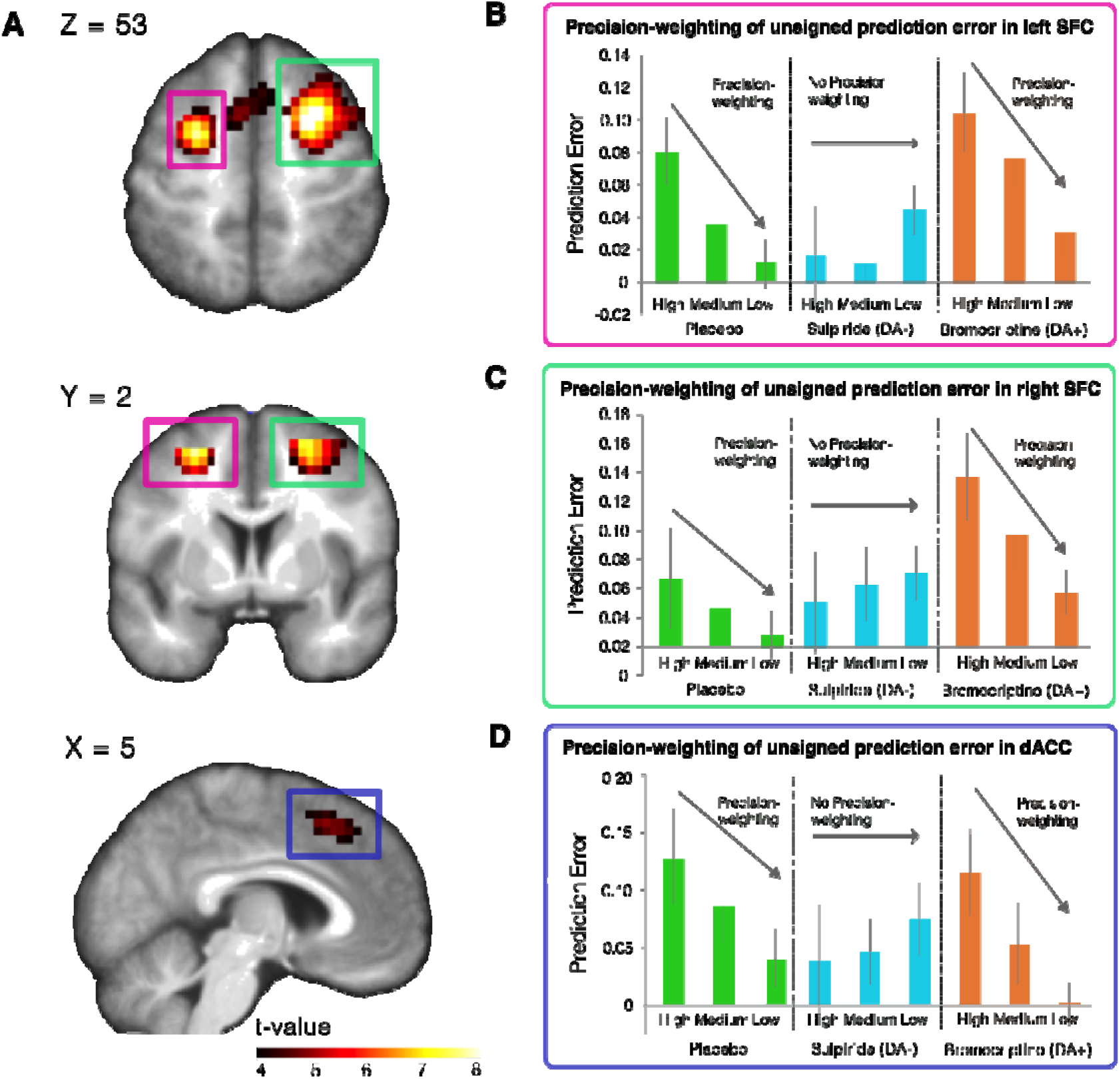
A. Unsigned prediction errors were coded in bilateral superior frontal cortex and dorsal anterior cingulate cortex. The left side of the brain is the left side of the image. B-D. When exploring these regions further, we find that unsigned prediction errors are coded in a precision-weighted fashion as indicated by the strong unsigned prediction error signal in the high precision condition which declines over the medium and low precision condition in the placebo and bromocriptine group. Importantly, sulpiride perturbed precision-weighting significantly in the left SFC. Error bars represent standard error of the mean.

#### Precision-weighting of unsigned prediction errors is mediated by dopamine in the SFC/dACC

To test whether precision and dopaminergic perturbations affected the coding of unsigned prediction errors, we extracted the parameter estimates (betas) of the unsigned prediction error parametric modulators from the left and right superior frontal cortex (SFC) and dACC cluster that showed a main effect of unsigned prediction error coding at whole brain corrected *p*FWE<.01. We used a two-factor mixed model ANOVA with medication group as the between-subjects variable and precision condition as the within-subjects variable, using a linear contrast across precision conditions for the main effect of precision and interaction. In the left SFC cluster, there was a significant interaction across precision conditions and medication group, suggesting that medication had a significant effect on precision-weighting of unsigned prediction errors (F{2,56}=4.025 *p*=.023; Fig.3B). There was a significant interaction between medication group (placebo vs sulpiride) and precision condition (F{2,37} = 5.44, *p*=.025), with less precision-weighting in the sulpiride than in the placebo group, which suggests that sulpiride dampens precision-weighting of unsigned prediction errors. Comparing the placebo and bromocriptine group, there was a significant effect of precision (F{1,37} = 14.93, *p*<.001), but no significant effect of medication group (placebo vs bromocriptine) (F{1,37} = 2.781, *p*=.104) or interaction between medication group and precision (F{2,36} = 0.02, *p*=.894). This finding suggests that left SFC unsigned prediction error signals are precision-weighted, but relatively unaffected by bromocriptine.

In the right SFC we did not find a significant interaction between medication and precision condition (F{2,56}=1.70, *p*=.193; Fig. 3C). However, signal changes in the right SFC are largely the same as in the left SFC (see Fig. 3B). We did find a significant main effect of medication (F{2,56}=3.65, *p*=.032). This effect was driven by a stronger main effect of unsigned prediction error in the bromocriptine group compared to the placebo group (F{1,37}=6.740, *p*=.013), whereas the difference was only trend-level significant between the sulpiride and placebo group (F{1,38}=3.55, *p*=.067).

In the dACC we found a trend-level significant interaction between precision and medication (F{2,56}=2.81, *p*=.069; Fig. 3D). Post-hoc tests between the placebo and sulpiride group revealed a trend-level interaction between precision and medication (F{1,38}=3.043, *p*=.089). Testing the placebo and bromocriptine group revealed a significant effect of precision (F{1,37}=9.32, *p*=.004), but no effect of group (F{1,37}=1.17, *p*=.29) or interaction (F{1,37}=.172, *p*=.68). A similar pattern was thus found in the dACC as in the left SFC (see Fig. 3B+D). The degree of cortical precision-weighting correlated with task performance (controlling for group), such that higher precision-weighting relates to better performance (Figure 4. Left Rho=-.45, *p*<.001; Right: Rho=-.40, *p*=.002; dACC: Rho=-.25, *p*=.055). There were no whole brain effects of group on precision-weighting.

**Figure 4:**
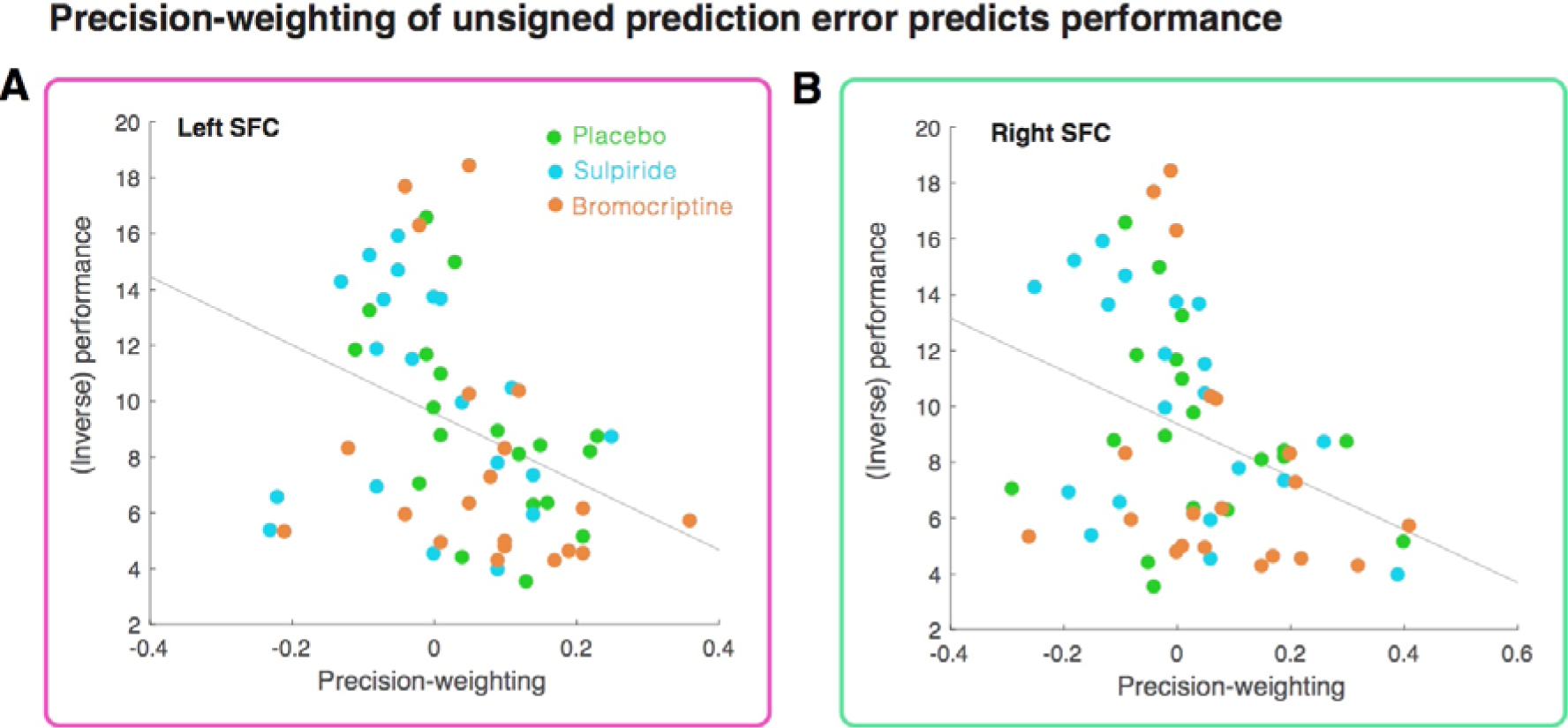
Precision-weighting of unsigned prediction errors in the left SFC (A) and right SFC (B) correlates with performance (i.e. difference between mean of the reward distribution and predicted mean) on the task.

### Study 2: Psychosis study

#### First Episode Psychosis (FEP) is associated with decreased overall performance and less benefit from more precise information

We explored the difference between participants’ estimates of the mean and the actual mean across trials, and tested for significant differences across groups and precision conditions while correction for multiple comparisons using a Bonferonni correction. We found better performance when precision was high in healthy control participants and ARMS individuals but not in people with FEP. We also found decreased performance in the FEP group compared to controls. There was a trend-level difference in number of missed trials, suggesting that on average the healthy controls missed one trial less then the other groups. There were no other significant differences in RT and scrolling distance (see Fig. 5 and Supplementary results).

**Figure 5:**
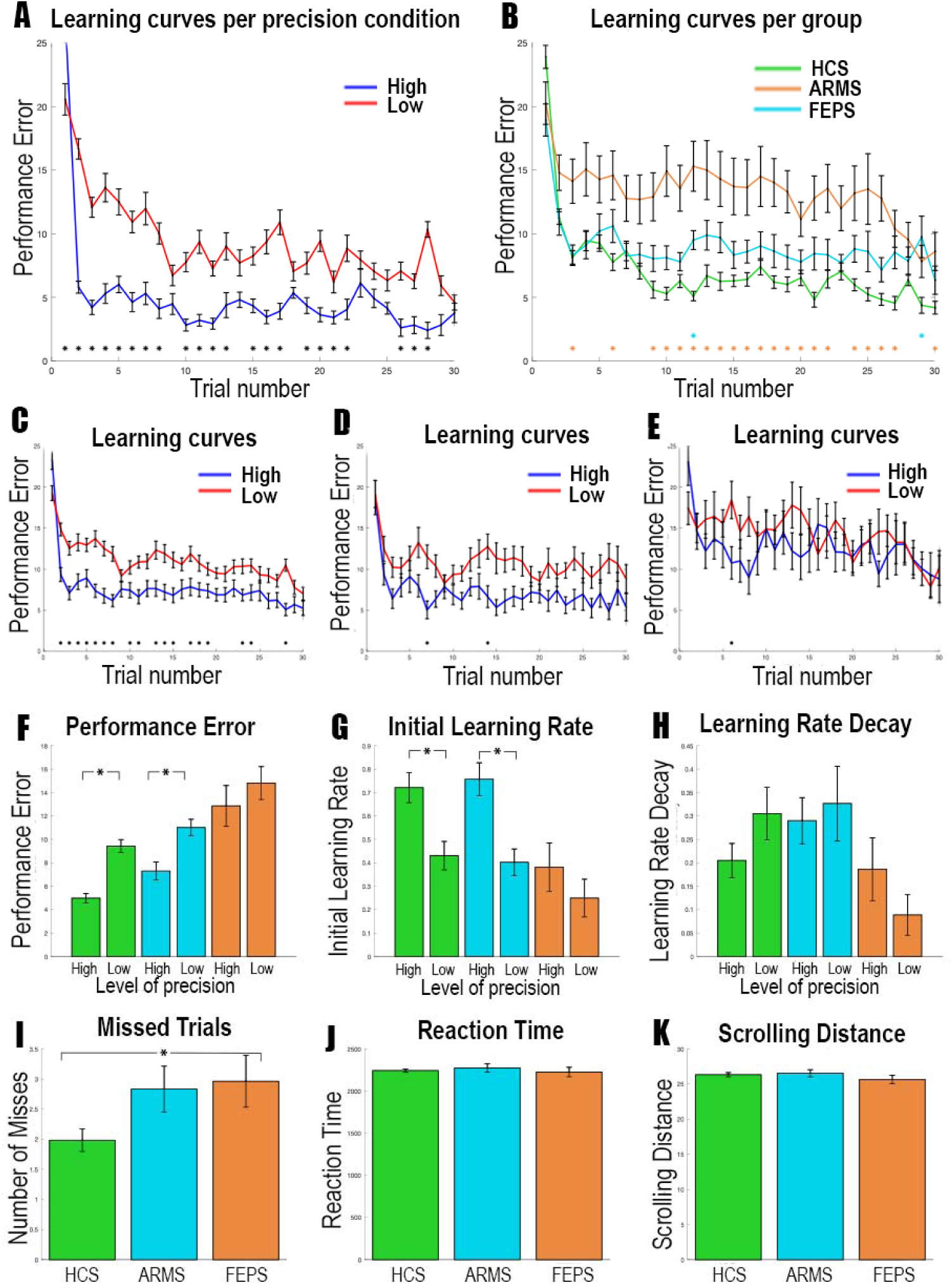
Behavioural results for the patient study; green bars are healthy controls, blue ARMS, and orange first episode psychosis. A-E display the average learning curves, reflecting the absolute distance between the actual mean of the distribution and the participants estimate of the mean distribution over 30 trials averaged over the 3 sessions for each participant. The distance between the prediction and the actual mean of the distribution defines performance error. Therefore, lower values of performance are better. Asterisks indicate Bonferroni corrected significant differences across conditions. A: Performance was significantly better in the high precision condition compared to the low precision condition when combing all participants, especially at the beginning of the experiment. B: HC performed better than FEPs, but not compared to ARMS. The colour of the asterisk indicates a significant difference of the patient group with HC. C-E: Healthy controls (C) showed a significant difference between the precision conditions, whereas the patient groups did not (D= ARMS, E= FEP). F: Averaging performance error across all trials, HC and ARMS perform better than FEP, and benefitted from more precise information, whereas FEP did not. G: Learning rates were higher in the high precision condition compared to the low precision condition in HC and ARMS, but not for FEPS, although an interaction was not significant. H: Precision did not affect the learning rate decay parameter. I: HC had slightly fewer missed trials than FEP. J: Reaction times were equal across groups. K: Scrolling distance was equal across groups. Error bars represent standard error of the mean.

#### FEP is associated with a lack of precision-weighting as revealed by computational modelling

We found that for the HCS and ARMS participants the best model of behaviour was the Pearce-Hall model with a precision-weighting parameter for both the signed and unsigned prediction error term. However, the FEP group followed a simple RW-learning rule without precision-weighting of prediction error, suggesting that the FEP group specifically is not precision-weighting prediction errors (Supplementary Table 3). This is further supported by the observation that HCS and ARMS participants show higher learning rates in the high precision condition, whereas the FEP group does not (Figure 5G and Supplementary results). When we correlated participants’ behavioural response data with data simulated using the individual parameters of the winning model for each group, we found that there was no difference in the amount of variance explained between groups (ANOVA: F{2,79}=.72, p=.48, r-values: HC = .40, ARMS = .34, FEP = .37), suggesting that the modelling procedure was equally successful across groups (see Supplementary Figures 1-3).

#### Unsigned prediction-errors are precision-weighted bilaterally in the SFC

In bilateral SFC (see methods for ROI derivation) there were brain signals that encoded unsigned prediction error (Right: T=7.33, voxels: 150, *p*<.001, [24 4 52]; Left: T=6.78, voxels: 149, *p*<.001, [−24 6 52], small volume correction). There was significantly stronger encoding of unsigned prediction errors in the high precision condition compared to the low precision condition in both the right and left SFC, demonstrating precision-weighting (Right: T=3.82, voxels: 66, *p*=.011, [22 12 52]; Left: T=3.52, voxels: 45, *p*=.025, [−21 −2 52]; small volume corrected), which is consistent with the effect observed in the dopaminergic modulation study.

In a whole brain analysis, additional regions demonstrated precision weighting of prediction error: bilateral superior frontal cortex, right lateral frontal cortex, and medial parietal lobe (Supplementary Results).

We also tested for signed prediction errors in the ventral striatum and the midbrain using ROI’s based on Diederen et al., 2017. However, no significant voxels were found that coded a main effect of signed prediction errors, a precision-weighting effect or a precision by group interaction (all *P* >.1).

#### FEP is associated with diminished precision-weighting in the right SFC

There was a significant difference in precision-weighting of prediction error between the FEP group and the control group in the right SFC (T=3.38, voxels: 9, *p*=.035,[24 9 48]; small volume corrected) (see Fig. 6A). Importantly, group differences were not driven by medication as precision-weighting in medicated psychosis patients was not significantly different from patients who non-medicated (T{18}=.14, p=.89; analysis conducted on voxels that showed a FEP v control group difference), and there was no correlation between medication dose and precision-weighting (r=.17, p=.52). No voxels differentiated the groups on whole-brain analysis, or left SFC ROI analysis, corrected for multiple comparisons. We tested whether precision-weighting in these 9 voxels (that differentiated the first episode psychosis and control groups) correlated to positive symptom severity (sum of PANSS items P1,2&3). To increase the number of participants for this analysis with a wide variety of symptoms we included both the ARMS group the FEP group (Fig 6B). Reduced precision-weighting related to greater positive symptoms (r = -.33, *p* = .032), but not when controlling for group (*p*=.3). As group and symptoms are confounded given our FEP inclusion criterion of having current delusions and/or hallucinations, and as low sample size limits our statistical power for correlations within group, we also ran an additional analysis including an extra 6 participants with FEP who had currently too low levels of positive psychotic symptoms to be included in main study (pooled ARMS & FEP, controlling for group r=-0.28, p=0.054; see Supplementary material for more details).

**Figure 6:**
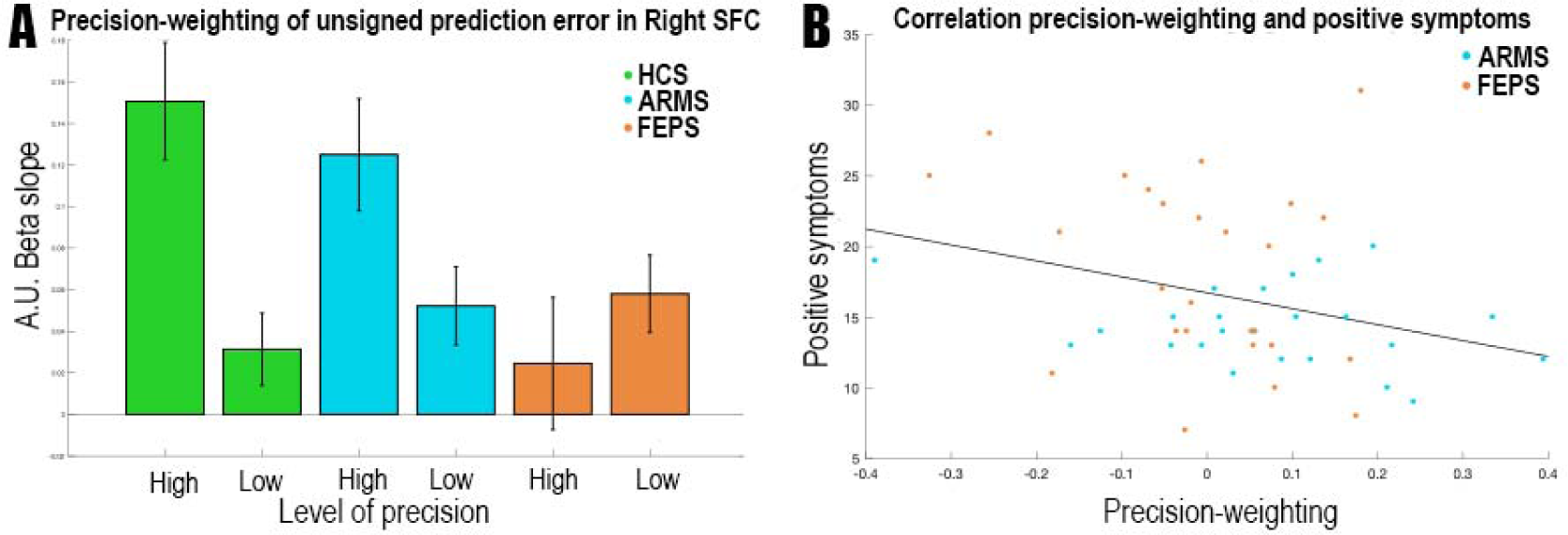
A: Precision-weighting of prediction error in Superior Frontal Cortex region of interest. Precision-weighting is significantly diminished in first episode psychosis. The y-axis provide beta estimates for the unsigned prediction error in different precision conditions in arbitrary units (a.u.). B: Diminished precision-weighting of prediction error is correlated with positive symptoms. Error bars represent standard error of the mean.

#### Higher schizotypy is related to decreased performance and diminished precision-weighting of cortical prediction errors in a separate healthy sample

We next examined the relationship between schizotypy and precision-weighting of prediction error in health, free from the possible confounds of medication or illness duration driving effects (Supplementary methods). We pooled the participants in the dopaminergic modulation study (which is the study described in this paper, N=59) and the participants of a previously collected healthy sample (who are from a previously reported study (Diederen et al., 2016), N=27) and tested for a relationship between schizotypy and precision-weighting of prediction error, while controlling for experimental group. There was a significant correlation between performance and schizotypy (Rho=-.23, *p*=.034). This was mirrored by a significant brain signal-schizotypy correlation between schizotypy and the extracted right SFC precision-weighting parameter estimates (r=-.25, *p*=.024) (Figure 7). The higher schizotypal personality, the less the participants exhibited cortical precision-weighting of prediction errors. No relationship was observed between schizotypy and the main-effect of prediction error (*p*>.3), suggesting that the effect is specific to the precision-weighting of prediction error.

**Figure 7:**
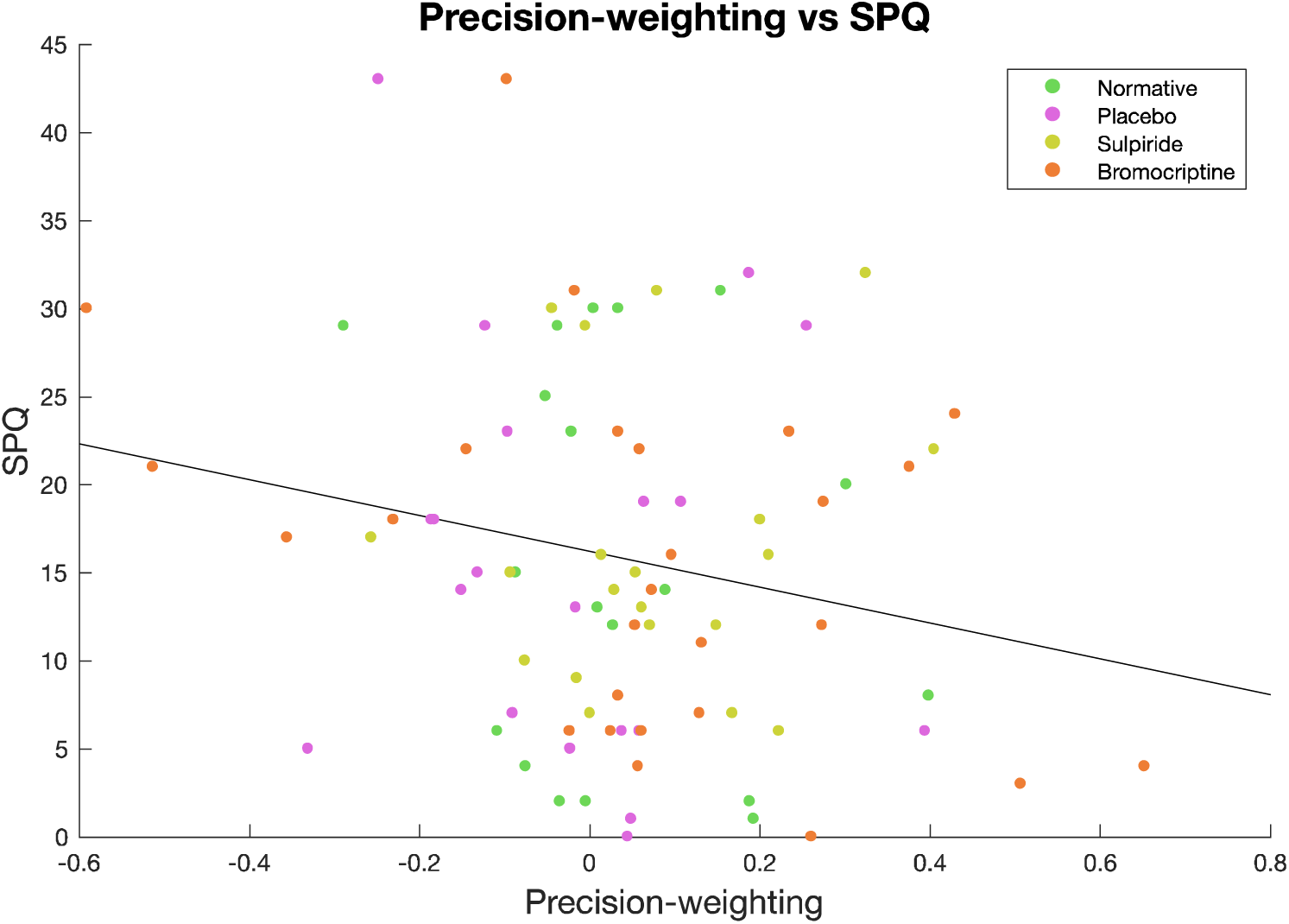
Higher SPQ scores (schizotypy) were correlated with less precision-weighting in the right Superior Frontal Cortex, including when controlling for experimental group; Rho=-.25, p=.024

## Discussion

We show that unsigned prediction errors are coded in superior frontal cortex, where the unsigned prediction error signal is coded relative to the precision of environmental outcomes; that the degree of precision-weighting benefits learning, is mediated by dopamine, is perturbed in first episode psychosis, and relates to schizotypy in a health.

Recent theories (Friston, 2009; Bastos et al., 2012; Adams et al., 2013; Fletcher & Frith, 2009; Sterzer et al., 2018) have hypothesised that precision-weighting of cortical prediction errors is mediated by neuromodulators (including dopamine), and link a malfunctioning dopamine system to psychosis through aberrant precision-weighting of these prediction errors. However, to our knowledge, no direct evidence for any of these claims exists. Here we showed that separately estimated precision-weighted signed and unsigned prediction errors provided the best description of the behavioural data, thus suggesting that both precision-weighted and unsigned prediction errors should be represented in the brain. The representation of precision weighted *signed* prediction errors in subcortical areas was confirmed previously by fMRI in humans (Diederen et al., 2016). In the present study, we tested the prediction that unsigned prediction errors would be represented in the brain, and found evidence for a precision weighted cortical representation of unsigned prediction in the bilateral superior frontal cortex and dorsal anterior cingulate cortex/pre-supplementary motor area (that we collectively referred to as SFC). Cortical precision weighting was significantly diminished in the sulpiride (dopamine D2 receptor antagonism) group in comparison to the other groups in the right SFC; there was marginal evidence of a medication effect in the dACC. This finding suggests that dopamine plays a key role in the mechanisms underlying precision-weighting of unsigned prediction errors. We furthermore found that a greater degree of superior frontal precision-weighting of unsigned prediction error was significantly correlated to performance on the task, where an increase in precision-weighting resulted in more accurate predictions of upcoming rewards. These results confirm the prediction that there exist cortical unsigned prediction error signals, which influence performance and are precision weighted by dopamine.

The coding of unsigned prediction errors in the superior and middle frontal gyri and dACC is in line with earlier findings by Hayden (et al., 2011) who found unsigned prediction errors in the dACC of monkeys, and with prior fMRI studies in humans (Fletcher et al 2001, Turner et al 2004, Fouragnan et al., 2017; Fouragnan et al 2018; Metereau & Dreher 2012; Ide et al., 2013). Our findings are consistent with those of Katthagen et al (2018), who used reaction time data (rather than choice data) from a human fMRI reversal learning study to derive a relevance weighted unsigned prediction error signal, which was also represented in the dACC. Our data in the dopaminergic modulation study, replicated in the psychosis study, show (for the first time to our knowledge) that cortical prediction error signals based on choice data are precision-weighted in humans. We note that dopaminergic innervation of cortex is greatest in superior frontal regions (Lewis et al., 1987; Berger et al., 1991; Paus, 2001), compatible with the hypothesis that precision weighting is influenced here by dopaminergic input.

If the precision-weighting of prediction errors is important in learning, we can expect aberrant learning to occur when prediction errors are not scaled optimally to the environmental statistics determining the precision of available information. We tested whether this mechanism could be of importance to psychosis, which is characterized by delusional beliefs and hallucinatory perception (Fletcher & Frith et al., 2009). Previous work showed aberrant cortical and subcortical prediction error coding in people with psychosis (Murray et al., 2008; Corlett et al., 2007, Ermakova et al., 2018). As psychosis has consistently been associated with dopamine dysfunction (Howes & Kapur 2009), it is possible that a dopamine mediated precision-weighting process would be impaired in psychosis. Indeed, it has been suggested that dopamine dysregulation causes psychosis due to affecting the brains capacity to precision-weight prediction error (Adams et al., 2013). That is, if unreliable prediction errors were given excessive weight, they could have an exaggerated influence on driving changes in the brain’s model of the world, thereby contributing to the formation of abnormal beliefs. We found several lines of evidence suggesting that FEP in particular is associated with a failure to precision-weight prediction errors. First, FEP was associated with decreased performance on the task. Furthermore, the FEP group did not benefit as much from more precision reward information than the healthy controls and ARMS group did, and computational modelling indicated that the FEP group does not precision-weight prediction errors, as they follow a simple RW learning rule without precision-weighted prediction errors. By contrast, controls and ARMS follow a Pearce-Hall learning rule with precision-weighted prediction errors. This invites the question whether the poor performance of the FEP group might have more to do with the failure to diminish their learning rate appropriately over time (as this is what characterizes a Pearce-Hall model). However, subsequent analysis revealed that whereas controls and ARMS show a clear effect of precision on learning rate, the FEP group does not. In contrast, no differences were found for the decay parameter, suggesting that the differences lie in how much prediction errors are used in different precision conditions. Thirdly, neural evidence suggests that the FEP group does not precision-weight cortical prediction errors to the extent that healthy controls do, and that the degree of neural abnormality may relate to positive psychotic symptom severity (we acknowledge that the modest patient sample size and marginal significance of the correlation is not conclusive, though the relation is supported by the finding that in healthy individuals, the degree of cortical precision weighting relates to schizotypy, consistent with a continuum model of psychosis).

Several other studies have used this computational framework to study learning in individuals with psychosis that imply a failure to precision weight prediction errors. Powers (et al., 2017) used hierarchical Bayesian models to make inferences about the way individuals with psychosis respectively form beliefs about the environment. Critically in these models a prediction error is weighted by the precision of beliefs regarding cue-outcome contingencies, and the volatility of these relationships (Mathys et al., 2011). As such, these models imply precision-weighting of prediction, however they do not test the degree to which these prediction errors are precision-weighted explicitly. Our results complement these studies and provide an additional direct test of the degree of precision weighting of prediction errors in psychosis.

A previous study has reported differences between healthy controls and individuals with schizophrenia in the degree to which they adapt the coding of value to the variability in the environment (Kirschner et al., 2016). This process of adaptive coding is similar to precision weighting of unsigned prediction errors, as it reflects the brains capacity to scale neural signals to what is referred to as economic ‘risk’, in other words the spread of possible reward outcomes. In combination with the present findings, psychotic disorder might be associated with a broader failure to adapt neural signals to the statistics of the environment. We thus conclude that there is evidence for a diminishment in precision-weighting of unsigned prediction errors in individuals with first episode psychosis. This was most strongly related to the intensity of the positive symptoms experienced by the patients in this study. Our current study provides evidence for a key hypothesis in the field of predictive coding theories of psychosis, which is that psychosis is associated with a failure to accurately take into account the reliability of new information, leading to the formation of aberrant inferences about the world, predisposing to delusional beliefs. The finding that the degree of precision weighting of cortical prediction errors is modulated by dopamine, combined with the finding of abnormal precision-weighting in psychosis, is consistent with the posit that the origins of the precision-weighting deficit in psychosis are dopaminergic. However, we note that although we demonstrate dopaminergic modulation of the degree of precision weighting, there may be other neurotransmitters that also contribute to this process. As we did not measure dopamine function in the clinical studies, it remains possible that the patient deficits are secondary to non-dopaminergic mechanisms. Pharmacological fMRI in patients, or combined fMRI and PET studies (including dopaminergic ligands) in patients, would be required to test this.

In conclusion, we found evidence of precision-weighted unsigned prediction errors in the superior frontal and dorsal anterior cingulate cortices. Furthermore, we found that the precision-weighting of prediction errors was modulated by the dopaminergic antagonist sulpiride, and we found that the degree of precision-weighting in this area was correlated to performance on the task, providing evidence for the first time that dopamine plays a role in precision-weighting of unsigned prediction error brain signals during learning. Healthy people, but not patients with first episode psychosis, take into account the precision of the environment and unsigned prediction errors when updating beliefs; accordingly, the cortical unsigned prediction error signal is abnormal in psychotic illness, and relates to trait levels of schizotypy in the healthy population, implicating it as a key mechanism underlying the pathogenesis of psychotic symptoms.

## Acknowledgements

We thank the staff of the Clinical Research Facility and Wolfson Brain Imaging Centre, Cambridge Biomedical Research Centre, Rachel Anderson and Eleanor van Sprang for their help in data collection, CAMEO staff for help with recruitment, the participants, Martin Vestergaard for his help with the original modelling scripts that the models used here build on, and thank Wolfram Schultz for his role in designing the experimental paradigm and pharmacological study.

## Author roles

J.H.: Conceptualization, methodology, formal analysis, writing (original draft preparation, review and editing); P.C.F.: Conceptualization, project administration, funding acquisition, writing (review and editing). J.D.G Conceptualization, investigation, methodology, project administration. H.T. project administration, implementation, formal analysis. T.S. and H.Z. investigation, project administration, writing (review and editing). I.G. Conceptualization, investigation, methodology, project administration. K.M.J.D. Conceptualization, methodology, project administration, implementation, formal analysis, supervision, writing (original draft preparation, review and editing) G.K.M., conceptualization, project administration, methodology, supervision, writing (original draft preparation, review and editing).

## References

Adams, R. A., Stephan, K. E., Brown, H. R., Frith, C. D., & Friston, K. J. (2013). The computational anatomy of psychosis. Frontiers in Psychiatry, 4(May), 47.

Ashburner, J., & Friston, K. J. (2005). Unified segmentation. Neuroimage, 26(3), 839–851.

Bastos, A. M., Usrey, W. M., Adams, R. A., Mangun, G. R., Fries, P., & Friston, K. J. (2012). Canonical microcircuits for predictive coding. Neuron, 76(4), 695–711.

Berger, B., Gaspar, P., & Verney, C. (1991). Dopaminergic innervation of the cerebral cortex: unexpected differences between rodents and primates. Trends in neurosciences, 14(1), 21–27.

Clark, A. (2013). Whatever next? Predictive brains, situated agents, and the future of cognitive science. Behavioral and Brain Sciences, 36(3), 181–204.

Clark, A. (2015). Surfing uncertainty: Prediction, action, and the embodied mind. Oxford University Press.

Courville, A. C., Daw, N. D., & Touretzky, D. S. (2006). Bayesian theories of conditioning in a changing world, 10(7).

Corlett, P. R., Murray, G. K., Honey, G. D., Aitken, M. R. F., Shanks, D. R., Robbins, T. W., … Fletcher, P. C. (2007). Disrupted prediction-error signal in psychosis: Evidence for an associative account of delusions. Brain, 130(9), 2387–2400.

Corlett, P. R., Honey, G. D., & Fletcher, P. C. (2007). From prediction error to psychosis: ketamine as a pharmacological model of delusions. Journal of Psychopharmacology (Oxford, England), 21(3), 238–252.

Davies, D. (2017). Psychotic experiences beyond psychotic disorders: from measurement to computational mechanisms. PhD Thesis. University of Cambridge.

D’ardenne, K., McClure, S. M., Nystrom, L. E., & Cohen, J. D. (2008). BOLD responses reflecting dopaminergic signals in the human ventral tegmental area. Science, 319(5867), 1264–1267.

Diederen, K. M. J., & Schultz, W. (2015). Scaling prediction errors to reward variability benefits error-driven learning in humans. Journal of Neurophysiology, 114(3), 1628–1640.

Diederen, K. M. M. J., Spencer, T., Vestergaard, M. D. D., Fletcher, P. C. C., & Schultz, W. (2016). Adaptive Prediction Error Coding in the Human Midbrain and Striatum Facilitates Behavioral Adaptation and Learning Efficiency. Neuron, 90(5), 1127–1138.

Diederen, K. M. J., Ziauddeen, H., Vestergaard, M. D., Spencer, T., Schultz, W., & Fletcher, P. C. (2017). Dopamine Modulates Adaptive Prediction Error Coding in the Human Midbrain and Striatum. Journal of Neuroscience, 37(7), 1708–1720.

Ermakova AO, Knolle F, Justicia A, Bullmore ET, Jones PB, Robbins TW, Fletcher PC, Murray GK (2018). Abnormal reward prediction-error signalling in antipsychotic naive individuals with first-episode psychosis or clinical risk for psychosis. Neuropsychopharmacology. 43(8):1691–1699.

Fletcher, P. C., Anderson, J. M., Shanks, D. R., Honey, R., Carpenter, T. A., Donovan, T., & Bullmore, E. T. (2001). Responses of human frontal cortex to surprising events are predicted by formal associative learning theory. Nature neuroscience, 4(10), 1043.

Fletcher, P. C., & Frith, C. D. (2009). Perceiving is believing: a Bayesian approach to explaining the positive symptoms of schizophrenia. Nature Reviews. Neuroscience, 10(1), 48–58.

Fouragnan, E., Queirazza, F., Retzler, C., Mullinger, K. J., & Philiastides, M. G. (2017). Spatiotemporal neural characterization of prediction error valence and surprise during reward learning in humans, Scientific Reports 7: 4762

Fouragnan, E., Retzler, C., & Philiastides, M. G. (2018). Separate neural representations of prediction error valence and surprise: Evidence from an fMRI meta-analysis. Human brain mapping. 39(7):2887–2906.

Friston, K. (2009). The free-energy principle: a rough guide to the brain?. Trends in cognitive sciences, 13(7), 293–301.

Friston, K. J., Stephan, K. E., Montague, R., & Dolan, R. J. (2014). Computational psychiatry: The brain as a phantastic organ. The Lancet Psychiatry.

Friston, K., Schwartenbeck, P., FitzGerald, T., Moutoussis, M., Behrens, T., & Dolan, R. J. (2014). The anatomy of choice: dopamine and decision-making. Phil. Trans. R. Soc. B, 369(1655), 20130481.

Gershman SJ. (2015). A Unifying Probabilistic View of Associative Learning. PLoS Comput Biol. 11(11):e1004567.

Hall, G., & Pearce, J. M. (1982). Restoring the associability of a pre-exposed CS by a surprising event. The Quarterly Journal of Experimental Psychology Section B, 34(3b), 127–140.

Hayden, B. Y., Heilbronner, S. R., Pearson, J. M., & Platt, M. L. (2011). Surprise Signals in Anterior Cingulate Cortex⍰: Neuronal Encoding of Unsigned Reward Prediction Errors Driving Adjustment in Behavior, 31(11), 4178–4187.

Heinz, A., Murray, G.K., Schlagenhauf, F., Sterzer, P., Grace, A.A., Waltz, J.A., (2018) Towards a Unifying Cognitive and Computational Neuroscience Account Of Psychotic Experience. Schizophrenia Bulletin doi: 10.1093/schbul/sby154.

Hohwy, J. (2013). The predictive mind. Oxford University Press.

Howes, O. D., & Kapur, S. (2009). The dopamine hypothesis of schizophrenia: version III—the final common pathway. Schizophrenia bulletin, 35(3), 549–562.

Ide, J. S., Shenoy, P., Yu, A. J., & Li, C. R. (2013). Bayesian Prediction and Evaluation in the Anterior Cingulate Cortex, 33(5), 2039–2047.

Katthagen, T., Mathys, C., Deserno, L., Walter, H., Kathmann, N., Heinz, A., & Schlagenhauf, F. (2018). Modeling subjective relevance in schizophrenia and its relation to aberrant salience. PLoS Computational Biology, 14(8), e1006319.

Kirschner, M., Hager, O. M., Bischof, M., Hartmann-Riemer, M. N., Kluge, A., Seifritz, E., & Kaiser, S. (2016). Deficits in context-dependent adaptive coding of reward in schizophrenia. NPJ schizophrenia, 2, 16020.

Lewis, D. A., Campbell, M. J., Foote, S. L., Goldstein, M., & Morrison, J. H. (1987). The distribution of tyrosine hydroxylase-immunoreactive fibers in primate neocortex is widespread but regionally specific. Journal of Neuroscience, 7(1), 279–290.

Mathys, C., Daunizeau, J., Friston, K. J., & Stephan, K. E. (2011). A Bayesian foundation for individual learning under uncertainty. Frontiers in human neuroscience, 5, 39.

Metereau, E., & Dreher, J. C. (2012). Cerebral correlates of salient prediction error for different rewards and punishments. Cerebral Cortex, 23(2), 477–487.

Murray, G. K., Corlett, P. R., Clark, L., Pessiglione, M., Blackwell, a D., Honey, G., … Fletcher, P. C. (2008). Substantia nigra/ventral tegmental reward prediction error disruption in psychosis. Molecular Psychiatry, 13(3), 239, 267–276.

O’Doherty, J. P., Dayan, P., Friston, K., Critchley, H., & Dolan, R. J. (2003). Temporal difference models and reward-related learning in the human brain. Neuron, 38(2), 329–337.

O’Doherty, J., Dayan, P., Schultz, J., Deichmann, R., Friston, K., & Dolan, R. J. (2004). Dissociable roles of ventral and dorsal striatum in instrumental conditioning. science, 304(5669), 452–454.

Paus, T. (2001). Primate anterior cingulate cortex: where motor control, drive and cognition interface. Nature reviews neuroscience, 2(6).

Pearce, J. M., & Hall, G. (1980). A model for Pavlovian learning: variations in the effectiveness of conditioned but not of unconditioned stimuli. Psychological review, 87(6), 532.

Pessiglione, M., Seymour, B., Flandin, G., Dolan, R. J., & Frith, C. D. (2006). Dopamine-dependent prediction errors underpin reward-seeking behaviour in humans. Nature, 442(7106), 1042.

Powers, A. R., Mathys, C., & Corlett, P. R. (2017). Pavlovian conditioning–induced hallucinations result from overweighting of perceptual priors. Science, 357(6351), 596–600.

Rao, R. P. N., & Ballard, D. H. (1999). Predictive coding in the visual cortex: a functional interpretation of some extra-classical receptive-field effects. Nature Neuroscience, 2(1), 79–87.

Rescorla, R. A., & Wagner, A. R. (1972). A theory of Pavlovian conditioning: Variations in the effectiveness of reinforcement and nonreinforcement. Classical conditioning II: Current research and theory, 2, 64–99.

Schlagenhauf F, Huys QJ, Deserno L, Rapp MA, Beck A, Heinze HJ, Dolan R, Heinz A. (2014) Striatal dysfunction during reversal learning in unmedicated schizophrenia patients. Neuroimage. 89:171–80

Schultz, W., Dayan, P., & Montague, P. R. (1997). A neural substrate of prediction and reward. Science, 275(5306), 1593–1599.

Sterzer, P., Adams, R.A., Fletcher, P., Frith, C., Lawrie, S.M., Muckli, L., Petrovic, P., Uhlhaas, P., Voss, M. and Corlett, P.R. (2018). The predictive coding account of psychosis. Biological Psychiatry. 84(9):634–643.

Sutton, R. S., & Barto, A. G. (1998). Reinforcement learning: An introduction (Vol. 1, No. 1). Cambridge: MIT press.

Tian, J., Huang, R., Cohen, J. Y., Osakada, F., Kobak, D., Machens, C. K., & Watabe-Uchida, M. (2016). Distributed and mixed information in monosynaptic inputs to dopamine neurons. Neuron, 91(6), 1374–1389.

Turner, D. C., Aitken, M. R., Shanks, D. R., Sahakian, B. J., Robbins, T. W., Schwarzbauer, C., & Fletcher, P. C. (2004). The role of the lateral frontal cortex in causal associative learning: exploring preventative and super-learning. Cerebral Cortex, 14(8), 872–880.

Yung, A. R., Yuen, H. P., Phillips, L. J., Francey, S., & McGorry, P. D. (2003). Mapping the onset of psychosis: The comprehensive assessment of at-risk mental states (CAARMS). Schizophrenia Research, 60(1), 30–31.

