## Supplementary Material for "Precision-weighting of cortical unsigned prediction error signals benefits learning, is mediated by dopamine, and is impaired in psychosis"

*^1^*Department of Psychiatry, University of Cambridge, United Kingdom, *^2^*Wellcome Trust MRC Institute of Metabolic Science, Cambridge Biomedical Campus, United Kingdom, *^3^*Cambridgeshire and Peterborough NHS Trust, *^4^*Department of Psychosis studies, Institute of Psychiatry, Psychology and Neuroscience King's College London, United Kingdom, *^5^*Charité - Universitätsmedizin Berlin, Germany

***= joint last author/equal contributions**

**Correspondence to:**

Dr Diederen, Department of Psychosis studies, Institute of Psychiatry, Psychology and Neuroscience King's College London, De Crespigny Park, London, SE5 8AF, United Kingdom **or**

Dr Murray, Department of Psychiatry, University of Cambridge, Douglas House, Trumpington Road 18b, Cambridge, CB2 8AH, United Kingdom

**Supplementary methods**

Participants received a single oral dose of bromocriptine 2.5 mg (dopamine D2 agonist), sulpiride 600 mg (D2 antagonist), or placebo. Domperidone 10mg was added to prevent potential nausea, as it would be an indicative factor of having taken a drug, and could thus influence the blinding procedure, as well as making people feel unwell (see Diederen et al., 2017 for a detailed description of the dopaminergic manipulation, which reports an analysis focussed on a different psychological process and a different brain metric to the current study as well as having a different anatomical focus). fMRI scans were acquired 2.5 h after dosing to capture the window of maximal drug effect.

*Participants and intervention - dopamine study*

63 healthy volunteers were recruited for a pharmacology study exploring adaptation of reward prediction errors to context variability. 4 were excluded due to feeling unwell during scanning and/or a failure to complete the paradigm, or side-effects of the medication. All participants were recruited via the distribution of flyers in Cambridge and advertisements on the internet (Gumtree). Drug screenings were all negative. After receiving detailed information about the study, all participants gave written informed consent. Schizotypy (trait levels of schizophrenia-like thoughts and behaviour that occur on a continuum in the general population) was measured on these participants using the Schizotypal Personality Questionnaire SPQ (Raine et al., 1991). Prior to scanning, participants received a single dose of the D2-antagonist sulpiride (600mg), the dopamine agonist Bromocriptine (2.5 mg), or placebo, in a double-blind fashion. An additional 28 healthy participants (26 men; mean age 22.6 years; SD 5.4) were pooled with the participants of the dopamine study for the schizotypy brain-behaviour study (one excluded due to scanning issues); these participants were studied without any drug intervention.

*Task timings*

Each trial started with a fixation cross that was presented between 2100 and 4200ms. The fixation cross was followed by a cue that was presented for 500ms. Participants had 3500ms to make their prediction. Making the prediction involved scrolling a mouse ball with their fingers up and down, and clicking the left mouse button to state their prediction. This resulted in moving a bar up and down the screen on a scale that ranged from 0 to 100. The starting point on the bar was randomized throughout the experiment, to ensure that scrolling distance did not correlate with participants’ predictions. The participants were instructed to minimize the prediction error, i.e. the difference between the expected and the actual obtained reward. After the prediction, another fixation cross was presented for duration between 2100 and 5250ms. Thereafter, the obtained reward was presented in addition to the prediction and the reward prediction error for 1000ms (see Fig. 1 for an example trial).

*Pay-off*

If the participants would always receive the amount of money drawn by the computer, this might have reduced their motivation to reveal their true prediction. Therefore, the participants were rewarded according to their accuracy (i.e., proximity to the mean of the reward distribution) on 20% of the trials in order to keep them motivated to predict as well as possible.

*Behavioural analyses*

Behavioural data was analysed in Matlab. In order to test the performance of the participants, we analysed whether the predictions of the participants approached the mean of the distribution, during the experiment. We calculated the absolute difference between the mean of the distribution and the prediction in across all trials and averaged across the 3 sessions. We computed Bonferroni corrected paired t-tests to test the effect of precision across and within groups and the differences across groups (see figure 2&5A-E). We also tested for an effect of group and precision on average distance across the experiment. In addition, we analysed differences in scrolling distance to exclude the possibility that drug effects influenced willingness to do the task, resulting in a decrease in scrolling distance. We also analysed differences in reaction times and missed trials for the same reason.

In order to explore the contribution of signed and unsigned prediction error to learning we used a linear regression model with belief-updating as dependent variable and both unsigned and signed prediction error as independent variables. Belief updating here is defined as the change in value estimate from t_n_ to t_n+1_. That is, when the participant expects a value of 20 on t_1_ and expect as value of 30 on t_2_ there was a belief update of 10, which is expected to be explained by a prediction error experienced at t_1_. Since signed prediction errors, but not unsigned prediction errors inform the participant about the direction of learning, we expect the former but not the latter to have a main effect on belief-updating. Since formal learning models like the Pearce-Hall model (Pearce & Hall, 1980), suggest that unsigned prediction errors increase learning, we expect an interaction between unsigned and signed prediction error on belief-updating. In order to visualise the effect of signed and unsigned prediction errors on belief-updating, we created bins on the basis of signed prediction errors and plot belief-updating on the y-axis. If unsigned prediction errors interact with signed prediction error in predicting belief-updating we should expect to see a logit or sigmoid relationship between signed prediction error and belief updating.

*Behavioural computational modelling*

To investigate learning, we fitted several reinforcement learning models to participants’ prediction sequences. Each model used a common updating rule in which predictions on trial *n* depended on the prediction error and the learning rate on trial *n-1:*

**y_n_ = y_n-1_ + k_n_ δ_n_	Equation 1**

Here, y_n_ is the prediction made on trial *n*, *k_n_* refers to the learning rate and *δ_n­_* denotes the size of the prediction error*.* This is a standard reinforcement learning model, and allows to estimate to which degree prediction errors are being used by the participant by estimating the learning rate parameter (Sutton & Barto et al., 1998).

The first model consisted of a Rescorla-Wagner (RW) reinforcement learning model with a fixed learning rate. A fixed learning rate prescribes that each prediction error is weighted equally during learning (e.g., independent of whether a prediction error occurred at the start, or the end of a session):

**k_n_ = α	Equation 2**

The second model consisted of a Pearce-Hall (PH) model with a trial-wise, dynamic, learning rate, which prescribes higher weighting of prediction errors (i.e., more learning) at the start of a task session compared to later trials. In uncertain environments, it is more optimal to decrease the weighting of prediction errors as learning progresses (once participants become more certain of their predictions) as prediction errors will continue to occur as a result of the imposed uncertainty. The PH learning rate decays as trial progress, as a function of the previous learning rate and the experienced prediction error, which allows for fast updating of predictions at the beginning of each task session:

**k_n_ = ɣC|δ_n-1_| + (1 – ɣ) k_n-1_	Equation 3**

in which |δ| is the absolute prediction error, C is an arbitrary scaling coefficient and ɣ is the learning rate decay.

We additionally explored whether scaling prediction error to the reliability of the environment (i.e., precision–weighting) benefitted learning by comparing models that scaled the prediction error term and models that did not. We both explored models that simply precision-weighted the prediction error by a constant term ω (equation 4&5), and models that estimated the degree υ of precision-weighting for each individual (equation 6). The precision here is the true precision, and not the trial-by-trial estimated precision, as participants were informed prior to the experiment which of the cues indicated the high, medium and low variance condition. We estimated both models that had a single estimated υ for both signed and unsigned prediction errors and a model that had a separate variable for signed and unsigned prediction errors. In contrast to earlier studies (Diederen et al., 2015, 2016 & 2017) we used a linear precision-weighted model.

**y_n_ = y_n-1_ + k_n_ (δ/ ω)_n_	Equation 4**

**k_n_ = ɣC|(δ_n-1_/ ω)| + (1 – ɣ) k_n-1_	Equation 5**

**ω = 1-υ+ υ*(SD)	Equation 6**

For the Pearce-Hall model we explored the scaling of the signed and unsigned prediction error separately, as well as combined.

We fitted the free parameters to the subjective predictions by maximizing the log-likelihood. We used a combination of nonlinear optimization algorithms implemented in MATLAB to estimate the free parameters to each participant’s full dataset over the trials of all conditions.

We used the Akaike Information Criterion (AIC) to estimate the model evidence in favour of each of the two models. The model with the lowest AIC was used for further analyses.

*fMRI acquisition and pre-processing steps*

All pre-processing steps were performed using SPM8 (available at http://www.fil.ion.ucl.ac.uk; Wellcome Department of Cognitive Neurology, London, England) in combination with the Donders matlab (dmb) toolbox for combining data from different echo acquisition times. Data was acquired at 4 different echo times (TE’s) of 12, 27,91, 43,82 and 59,73 ms. Corrections for slice timing was not applied considering the relatively fast repetition times (TR) of 2100ms. Using SPM8 the functional images were realigned first, after which the four different echoes were summed and averaged in order to minimize signal intensity inhomogeneity due to differences in T2* relaxation times across the brain (Poser et al., 2006). The functional images were subsequently coregistered to the T1-weighted anatomical image in native-space by first coregistering one averaged functional image, after which the coregistration parameters of this registration were applied to the all the functional images in order to registered to native space. Unified segmentation was used in order to achieve the normalization of the functional and anatomical images to MNI space (Ashburner & Friston 2005). Unified segmentation is a single iterative model that combines segmentation, bias correction and normalization in order to achieve optimal results. The segmentation step was omitted for one of the participants; since it gave erroneous segmentation parameters resulting in distortions in the normalized anatomical T1 scan. The segmentation step was later conducted using a different template, namely the East-Asian template, which resulted in a correct segmentation. Spatial smoothing was performed using 8mm Gaussian kernels. The time series in each session were high-pas filtered at 128 Hz. These pre-processing steps were the same for all studies, with the exception that the patient study did not use a multi-echo sequence.

*2^nd^-level analysis dopamine study*

For the analysis of the main effect of unsigned prediction errors we first created 1^st^-level contrasts in which we combined the 2 EV’s for each separate precision condition, resulting in 3 contrasts corresponding to the 3 different precision conditions (high, medium, low). We modelled these 3 precision contrasts for each pharmacological condition separately specifying the effects to be dependent, and having equal variance. We subsequently tested for a main effect of prediction error by doing a positive 2^nd^-level contrast on the parametric modulators for the unsigned prediction error.

To test for an effect of precision on the coding of unsigned prediction error and the effect of pharmacological group, we extracted the parameter estimates (beta’s) for all precision conditions for each medication group separately from the left and right superior frontal cortex clusters and dACC cluster surviving FWE<.01 coding unsigned prediction errors; for each cluster we used the mean values per cluster per participants in a mixed-2-factor-ANOVA with a 3-level within subject factor (high, medium, low) and a 3-level between subject factor (medication), and tested for main effect of precision and interaction between precision conditions and medication using a mixed-model ANOVA in SPSS (version 21). Specifically we used a linear contrast (termed “linear polynomial contrast” by SPSS) across precision to examine the main effect of precision and interaction between group and precision. When a significant interaction was found we explored this further to see which group was driving the interaction.

*2^nd^-level analysis psychosis study*

For the analysis of the main effect of unsigned prediction errors we first created 1^st^-level contrasts in which we combined the 3 EV’s for each separate precision condition, resulting in 2 contrasts corresponding to the 2 different precision conditions (high, low). We modelled these 2 precision contrasts for each pharmacological condition separately specifying the effects to be dependent, and having equal variance. We subsequently tested for a main effect of prediction error by doing a positive 2^nd^-level contrast on the parametric modulators for the unsigned prediction error. We tested for a main effect of precision by doing a 2^nd^-level contrast on the two precision conditions. A small volume correction was implemented as follows: these tests were performed wholebrain uncorrected <.005, after which we used an ROI on the basis of the peak coordinates of the dopamine study [27 8 54] in form of a 10mm sphere, which was used to do a small-volume correction. We subsequently tested for differences in precision-weighting by creating a 1^st^-level contrast which subtracted the low-precision condition from the high-precision condition as a measure of precision-weighting. On 2^nd^-level we contrasted precision-weighting in the ARMS and FEP group against the HC group to test for significant differences in precision-weighting, again at <.005 uncorrected, after which we used the same ROI to test for differences in precision-weighting.

*2^nd^-level analysis schizotypy brain-behaviour study*

In order to test for a relationship between schizotypy and precision-weighting of prediction error, we extracted the betas from the precision-weighting contrast from a cluster which showed a significant main effect of unsigned prediction error at FWE<.01 in the dopamine study (which is the same criteria used for testing the effect of medication). In addition we re-analysed an existing data-set consisting of 27 healthy controls (pre-processing and 1^st^ level analysis is the same as in the dopamine study). We used an ROI (10mm sphere) based on the peak-voxel of the dopamine study [27 8 54], and tested for a main effect of prediction error at FWE<.01. Again we subtracted from these voxels the betas reflecting precision-weighting. Subsequently we used a general linear model to test for a relationship between SPQ scores and the extracted precision-weighting betas whilst controlling for medication group and study.

**Supplementary results: Study 1: Dopamine study**

*Environmental precision and D2 antagonism modulate task performance.*

First, we explored whether performance increases when environmental precision is high (i.e. when the standard deviation of rewarding outcomes is low). We measured performance by investigating how close participants’ predictions were to the actual mean of the distribution on average (i.e., the average difference between the mean of the reward distribution). We used a two-factor mixed model ANOVA with medication group as the between-subjects variable and precision conditions as the within-subjects variable, using a linear contrast across precision conditions for the main effect of precision and interaction between precision and medication. We found that performance increased when the precision increased (F{1,56}=11.3, *p*=.001), but there was no significant interaction between precision and medication group (F{2,56}=1.3, *p*=.27). However, there was a trend-level effect of group on average final performance (F{2,56}=2.5, *p*=.094). Post-hoc tests indicated reduced overall performance in the sulpiride group compared to placebo (F{1,38}=5.10, *p*=.030), but no significant interaction with precision (F{1,38}=.18, *p*=.68). Comparing the bromocriptine group to placebo there was no significant overall difference (F{1,37}=2.01, *p*=.17) and no significant interaction (F{1,37}=1.70, *p*=.20) (See Supplementary figure 1).

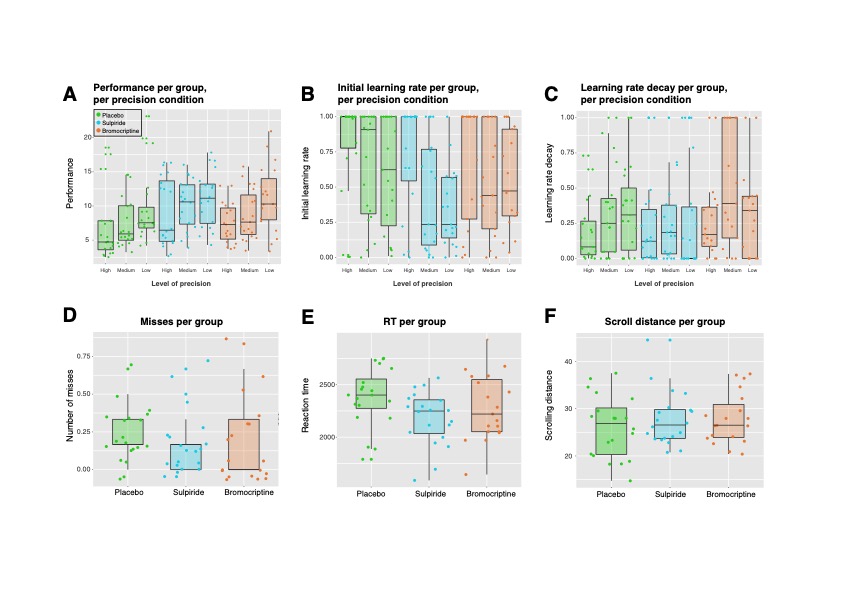

*Supplementary figure 1: Scatter box plots for performance stratified by precision condition and medication group (A), initial learning rates (B), learning rate decay (C), missed trials (D), average reaction time (E), and scrolling distance (F)*

*Elimination of possible confounding by effects on motivation*

Since dopamine also plays an important role in motivation, we explored whether the dopaminergic manipulation influenced measures of motivation, such as response times (Crespi, 1942; Niv, 2007), how much the participants moved the mouse ball to state their prediction (Fig. 3B; see methods section for more detail), and the number of missed trials (Fig. 3D). Using one-way ANOVAs, we found that only response times were significantly different across medication groups, with the sulpiride group being faster than placebo (RT: F{2,55}=4.21, *p*=.019; Misses: F{2,55}=0.33, *p*=.72; Scrolling distance: F{2,55}=0.56, *p*=.57; Fig. 3B-D). Differences in response times remained significant when controlling for scrolling distance: F{2,55}=3.32, *p*=.044. More rapid reactions in the sulpiride condition render it unlikely that any performance deficit secondary to sulpiride could be driven by motivational impairments. In the clinical psychosis study we also tested for differences in reaction times, scrolling distance, and missed trials, to see whether there could have been motivational and/ or motor/movement differences between groups. Using one-way ANOVA’s, we found that none of these measures were different across groups (RT: F{2,75}=1.77, p=.17; Misses: F{2,75}=0.1, p=.91; Scrolling distance: F{2,75}=0.12, p=.89) (Supplementary figure 1).

*Unsigned and signed prediction errors contribute to belief updating*

In order to explore whether signed and unsigned prediction errors contribute to learning, we fitted a simple regression model to the behavioural data, where the extent of learning, defined as the change in predicted reward from a trial to the next, was the dependent variable and signed and unsigned prediction errors were the independent variables. Signed prediction errors are defined in this task as the signed difference between predicted and received reward. Unsigned prediction errors are defined as the absolute difference between predicted and received reward. Both of these prediction errors can be observed directly in our task, and thus in contrast to many other experimental paradigms, do not have to be inferred as a latent variable using a modelling procedure. We expected a main effect of signed prediction error on learning, but not of unsigned prediction as the former informs the participants about the direction of belief updating, whereas the latter does not, and can therefore not drive learning directly. However, Pearce-Hall reinforcement learning models suggest that unsigned prediction errors increase the amount of attention devoted to a stimulus. Thus we expect that when unsigned prediction errors are high, the effect of signed prediction errors on learning is higher. That is, we expect there to be an interaction between unsigned and signed prediction errors. In line with this expectation, we found a significant interaction between signed and unsigned prediction errors (F{1,10763}=51.7, p<.0001), suggesting that learning is a function of both the signed and unsigned prediction error. In addition, we observed a significant effect of signed prediction errors on learning (F{1,10763}=246.95, p<.0001), whereas the effect of unsigned prediction errors was insignificant (F{1,10763}=.95, p=.33). We furthermore tested whether the precision condition interacted with signed prediction errors on belief updating. Although there was no main-effect of precision (F{2,10761}=.25, p=.78), there was indeed a significant interaction between precision and signed prediction error (F{2,10761}=25.8, p<.0001). The significant interaction between signed and unsigned prediction errors predicts a pseudo logit-function amplifying the contribution of signed prediction error on belief updating when the unsigned prediction errors are highest (i.e. when signed prediction errors are highly positive or highly negative). To visualise the interaction between signed and unsigned prediction error we binned signed prediction errors and plotted learning on the y-axis (Supplementary Figure 1A). We indeed find the predicted pseudo-logit shape between the relationship between signed prediction errors and learning such that the line is steeper when signed prediction errors are higher, indicating that unsigned prediction errors contribute to learning in addition to signed prediction errors. We furthermore visualised the effect of precision, where we see that signed prediction errors have a stronger effect on learning in the high precision condition compared to low precision condition (Supplementary Figure 1B). The significant interaction predicts a stronger effect of signed prediction error on belief updating in the high precision condition compared to the low precision condition. We thus conclude that signed prediction errors contribute more to learning when reward information is more precise. By simulating data using the estimated model parameters and correlating this with the acquired data, we show that the correlation between the simulated and the behavioural response data is comparable across groups (R^2^ placebo: .20 (SE= .035), sulpiride: .18 (SE= .034), bromocriptine: .19 (SE= .026) (ANOVA: *p*>.1), similar to other studies (e.g. Powers et al., 2017).

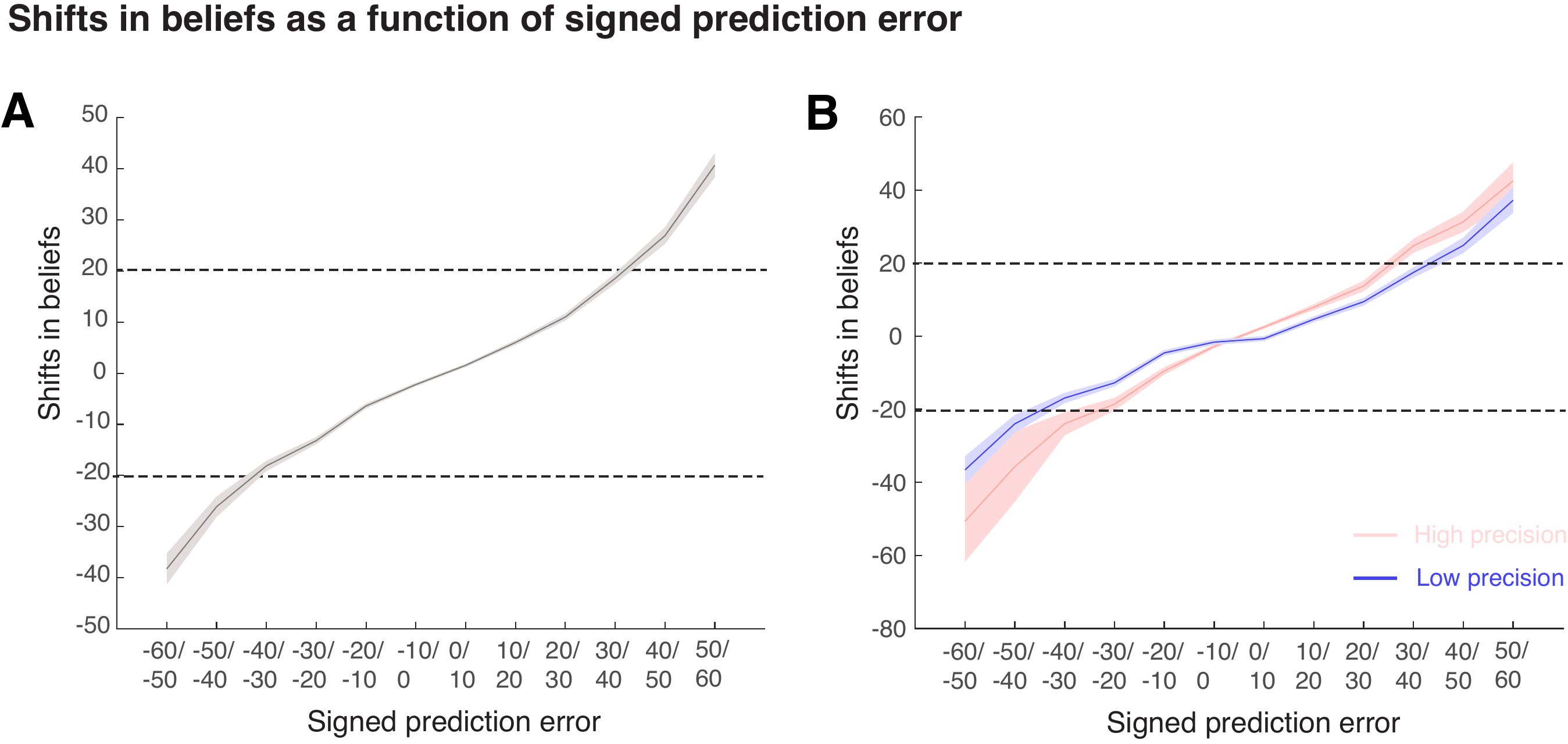

*Supplementary Figure 2: Here we show the relationship between signed prediction errors (x-axis) and belief-updating (y-axis) collapsed for the high, medium, and low precision conditions (A) and separate for high and low precision (B) (The medium precision condition is not plotted for clarity purposes). The shaded area is one standard error of the mean. In A the interaction effect between unsigned prediction error and signed prediction is shown (above the top and below the bottom dotted line unsigned prediction errors are strongest, amplifying the effect of the signed prediction error). In B the effect of precision is shown, revealing a stronger effect of signed prediction error in the high precision condition compared to low precision.*

*Bromocriptine increases learning rates*

The best-performing computational model contained 4 free parameters (initial learning rate, decay parameter, precision-weighting of signed prediction error and unsigned prediction error). As the model with a precision-weighting parameter outperformed the models without, precision is expected to affect the learning rate and decay parameters in the model. In order to explore this question further, we fitted the data to each precision condition separately for each participant and tested for an effect of precision on the model parameters. We tested for a main effect of group and precision on the initial learning rate and decay parameter (see methods), and their interactions. We found a significant effect of precision on the initial learning rate in the winning model, reflecting higher learning rates when reward information was more precise (F{1,52}=16.83, *p*<.001). In addition we found a small, but significant, effect of group (F{2,52}=3.252, *p*=.047), which was driven by higher learning rates in the bromocriptine group compared to the sulpiride group (F{1,36}=7.266, *p*=.011). However, no significant interaction was found F{2,52}=1.348, *p*=.27). We also explored the effects of precision and medication group on the decay parameter, and found no significant main effect of precision F{1,52}=2.48, *p*=.12), medication group F{2,52}=1.80, *p*=.18), or interaction F{2,52}=.44, *p*=.65) (see Fig. 2 G-H).

Supplementary Table 1: Overview of AIC values for each learning model per medication group. The lowest AIC, which indicates the best performing model, is printed in bold: a Pearce-Hall model with individual estimates of precision weighting for signed and unsigned prediction error. RW: Rescorla-Wagner. PH – Pearce-Hall.

|  | Free parameters | AIC overall | Placebo | Sulpiride | Bromocriptine |
| --- | --- | --- | --- | --- | --- |
| RW | 1 | 894.7 | 824.2 | 935.9 | 924.0 |
| RW scaled | 1 | 903.2 | 829.0 | 947.3 | 933.4 |
| PH not scaled | 2 | 884.2 | 808.2 | 931.1 | 913.4 |
| PH scaled | 2 | 883.6 | 807.3 | 930.1 | 913.4 |
| PH-estimated precision-weighting | 3 | 882.7 | 806.1 | 928.9 | 914.1 |
| **PH-estimated precision-weighting separate for signed and unsigned prediction error** | **4** | **879.4** | **803.1** | **926.3** | **910.4** |

Supplementary Table 2: Dopamine study. Regions coding unsigned prediction error voxelwise whole brain corrected <.05. SFC = Superior frontal cortex. SMA/dACC=supplementary motor area/dorsal anterior cingulate cortex. sOC= superior occipital cortex. FWE: family-wise error corrected.

| Region | MNI | Voxels | T | Z | *p*FWE |
| --- | --- | --- | --- | --- | --- |
| Right SFC | 27 8 54 | 145 | 8.43 | 7.68 | <.001 |
| Left SFC | -26 0 54 | 32 | 7.66 | 7.09 | <.001 |
| Pre SMA/ dACC | -7 12 50 | 6 | 5.32 | 5.10 | =.002 |
| Right sOC | 24 -74 34 | 21 | 7.24 | 6.75 | <.001 |
| Parietal cortex | 34 -48 38 | 18 | 5.69 | 5.43 | <.001 |
| Parietal cortex | 20 -60 26 | 4 | 5.24 | 5.03 | =.003 |

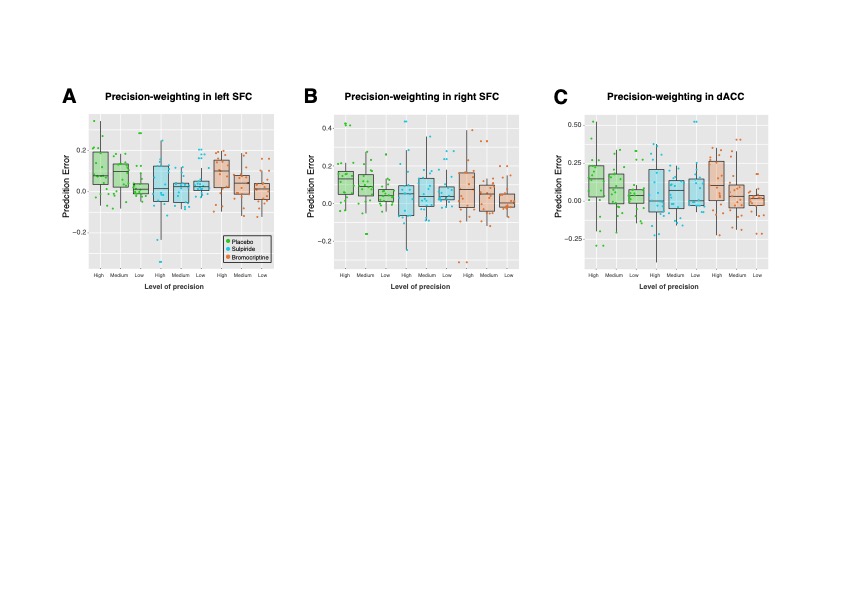

*Supplementary figure 3: Scatter boxplots for brain unsigned prediction errors signals across different precision conditions and groups in the left SFC(A), right SFC (B) and dACC (C). Sulpiride diminishes precision-weighting signals in these regions.*

*Secondary outcome variable analyses, dopamine study*

In addition to the analyses we conducted in our primary regions of interest, i.e. SFC and dACC, we also tested for an interaction between precision conditions and medication in the remainder of clusters that coded unsigned prediction errors. There were no significant precision by medication interactions: superior occipital cortex cluster, (F{2,56}=1.60, *p=*.21), larger parietal cluster (F{2,56}=.82, *p*=.45), smaller parietal cluster (F{2,56}=1.53, *p*=.23).

**Supplementary results: Study 2: Psychosis study**

*First Episode Psychosis (FEP) is associated with decreased overall performance and less benefit from more precise information*

First, we explored whether performance increases when environmental precision is high (i.e. when the standard deviation of rewarding outcomes is low). We measured performance by investigating how close participants’ predictions were to the actual mean of the distribution on average (i.e., the average difference between the mean of the reward distribution). We used a two-factor mixed model ANOVA with patient group as the between-subjects variable and precision conditions as the within-subjects variable, using a linear contrast across precision conditions for the main effect of precision and interaction between precision and medication. Performance improved when the precision increased (F{1,71}=81.4, *p*<.001), there was an effect of group (F{2,71}=14.9, *p*<.001) and there was a significant interaction between precision and patient group (F{2,71}=3.8, *p*=.028). Post-hoc tests comparing HCS to FEP indicated reduced overall performance in the FEP group compared to HCS (*p<.001*), and a significant interaction with precision (*p*=.018). ARMS was not significantly different from HCS (all *p*>.1). Still, the majority of participants in every group performed better (on inspection of average error stratified by condition) in the high precision condition (HCS: 29/30, ARMS: 20/24, FEP: 15/20).

Supplementary Table 3a: Overview model comparisons. RW: Rescorla-Wagner. PH – Pearce-Hall. FEP group is best fit by a simple Rescorla-Wagner Model. Other groups are best fit by a Pearce-Hall model with individual estimates of precision weighting for signed and unsigned prediction error. AIC is Akaike Information Criterion, BIC is Bayesian Information Criterion.

|  | Free parameters | AIC | | | | BIC | | | |
| --- | --- | --- | --- | --- | --- | --- | --- | --- | --- |
|  |  | Combined | HC | ARMS | FEP | Combined | HC | ARMS | FEP |
| RW | 1 | 1365.5 | 1316.1 | 1375.5 | **1077.1** | 1366.0 | 1319.3 | 1378.7 | **1407.9** |
| RW scaled | 1 | 1364.8 | 1311.7 | 1376.5 | 1077.3 | 1365.1 | 1314.9 | 1379.8 | 1409.4 |
| PH not scaled | 2 | 1359.8 | 1298.7 | 1362.0 | 1094.2 | 1363.0 | 1305.2 | 1368.4 | 1424.8 |
| PH scaled | 2 | 1361.4 | 1299.7 | 1362.9 | 1096.5 | 1364.6 | 1306.2 | 1369.3 | 1427.8 |
| PH-estimated precision | 3 | 1355.6 | 1292.9 | 1356.7 | 1097.1 | 1373.1 | 1317.2 | 1372.8 | 1437.8 |
| PH-estimated precision separate for signed and unsigned PE | 4 | **1349.8** | **1288.6** | **1348.5** | 1087.8 | **1359.5** | **1301.5** | **1361.4** | 1424.7 |

Table 3b. AIC values for the 6 models listed per participant

| HCS | RW | RW PW | PH | PH PW | PH est-PW | PH est-PW2 |
| --- | --- | --- | --- | --- | --- | --- |
| 1 | 1183.877504 | 1152.335075 | 1091.073479 | 1108.445366 | 1046.807444 | 1031.410302 |
| 2 | 1372.3567 | 1389.148124 | 1368.510955 | 1371.998807 | 1370.510955 | 1373.557244 |
| 3 | 1256.172648 | 1209.671898 | 1204.034971 | 1183.966858 | 1278.892349 | 1207.582963 |
| 4 | 1286.915137 | 1292.687063 | 1284.682754 | 1305.687781 | 1308.68253 | 1287.666483 |
| 5 | 1290.438873 | 1264.904237 | 1283.227608 | 1267.107432 | 1312.884255 | 1281.224949 |
| 6 | 1413.007476 | 1418.671756 | 1413.265389 | 1414.315255 | 1410.450647 | 1402.879401 |
| 7 | 1193.189563 | 1206.167926 | 1161.406215 | 1156.439688 | 1165.28439 | 1165.406215 |
| 8 | 1273.981587 | 1315.553137 | 1223.521654 | 1243.872301 | 1225.521654 | 1227.521654 |
| 9 | 1683.739862 | 1675.327576 | 1794.086382 | 1789.804444 | 1794.984204 | 1761.295382 |
| 10 | 1151.680045 | 1164.264302 | 1083.311238 | 1095.981541 | 1079.191284 | 1059.184916 |
| 11 | 1305.316798 | 1302.971452 | 1288.746089 | 1282.516355 | 1339.432822 | 1292.65664 |
| 12 | 1287.613281 | 1286.714175 | 1287.510228 | 1287.921629 | 1277.960161 | 1267.878723 |
| 13 | 1490.664035 | 1486.708187 | 1488.498582 | 1488.041862 | 1506.585259 | 1492.435485 |
| 14 | 1187.257141 | 1174.139719 | 1184.657132 | 1179.84371 | 1184.3263 | 1185.278296 |
| 15 | 1488.774125 | 1501.536743 | 1490.777241 | 1490.616645 | 1490.619594 | 1494.777241 |
| 16 | 1410.837226 | 1415.684792 | 1412.837226 | 1404.176203 | 1404.315796 | 1390.032547 |
| 17 | 1190.07592 | 1192.009307 | 1169.097198 | 1173.323817 | 1217.762025 | 1172.153952 |
| 18 | 1499.373384 | 1503.565039 | 1501.759155 | 1508.619215 | 1500.311065 | 1503.746359 |
| 19 | 1410.041057 | 1415.189265 | 1401.905387 | 1405.494594 | 1400.552992 | 1398.200362 |
| 20 | 1174.468765 | 1169.22675 | 1160.965826 | 1164.502561 | 1154.220974 | 1155.584923 |
| 21 | 1498.267087 | 1509.594791 | 1500.582459 | 1500.295521 | 1502.582459 | 1500.908157 |
| 22 | 1420.518099 | 1412.074755 | 1422.518099 | 1422.048845 | 1423.636762 | 1423.287055 |
| 23 | 1269.086796 | 1278.257371 | 1236.325299 | 1234.656857 | 1318.793 | 1240.325299 |
| 24 | 1318.006298 | 1337.070785 | 1305.853297 | 1296.587653 | 1298.717735 | 1322.494052 |
| 25 | 1144.815638 | 1060.319846 | 1086.486895 | 1041.108916 | 1078.417646 | 1071.931398 |
| 26 | 1259.25655 | 1231.916971 | 1208.445368 | 1218.388043 | 1152.90666 | 1129.550451 |
| 27 | 1318.715787 | 1300.411999 | 1293.599675 | 1293.32385 | 1314.442473 | 1296.4482 |
| 28 | 1206.507856 | 1198.292537 | 1160.122259 | 1208.750694 | 1265.708721 | 1145.12081 |
| 29 | 1347.226209 | 1337.872312 | 1340.619916 | 1338.081606 | 1318.946966 | 1304.251352 |
| 30 | 1150.802427 | 1147.263804 | 1113.882229 | 1115.646725 | 1082.486298 | 1073.190826 |

| ARMS | RW | RW PW | PH | PH PW | PH est-PW | PH est-PW2 |
| --- | --- | --- | --- | --- | --- | --- |
| 1 | 1294.061999 | 1294.998096 | 1276.635474 | 1267.525752 | 1277.299565 | 1277.747487 |
| 2 | 1380.415712 | 1402.652224 | 1382.415712 | 1378.631345 | 1384.415712 | 1372.988857 |
| 3 | 1477.633097 | 1490.27286 | 1472.683629 | 1478.976821 | 1460.311501 | 1454.399688 |
| 4 | 1397.324394 | 1396.015402 | 1398.491407 | 1394.818783 | 1393.795237 | 1383.155924 |
| 5 | 1396.659958 | 1402.366444 | 1398.659958 | 1399.110087 | 1400.631163 | 1405.001064 |
| 6 | 1473.826341 | 1470.523192 | 1475.826341 | 1474.407748 | 1476.277513 | 1463.979168 |
| 7 | 1353.187904 | 1381.199618 | 1355.187904 | 1365.686861 | 1357.187904 | 1359.187904 |
| 8 | 1402.86158 | 1419.654054 | 1404.86158 | 1404.883876 | 1406.86158 | 1408.86158 |
| 9 | 1518.669768 | 1518.470631 | 1510.895713 | 1511.737856 | 1511.576356 | 1512.919587 |
| 10 | 1139.540689 | 1131.504958 | 1087.974616 | 1096.547318 | 1046.267617 | 1035.980372 |
| 11 | 1445.309806 | 1459.306526 | 1456.260852 | 1457.623709 | 1455.942689 | 1451.268396 |
| 12 | 1191.608535 | 1184.932506 | 1164.798882 | 1166.561225 | 1166.73363 | 1167.824342 |
| 13 | 1419.053459 | 1407.149432 | 1421.053459 | 1415.578358 | 1418.121072 | 1390.296664 |
| 14 | 1434.076129 | 1438.021678 | 1408.994075 | 1414.82572 | 1410.992339 | 1399.660663 |
| 15 | 1271.308937 | 1272.882741 | 1257.185481 | 1250.986516 | 1256.856919 | 1255.63322 |
| 16 | 1547.133043 | 1547.616577 | 1560.113993 | 1558.348111 | 1561.987902 | 1562.959598 |
| 17 | 1464.432361 | 1476.657998 | 1479.423979 | 1472.715123 | 1488.777718 | 1466.605038 |
| 18 | 1527.363424 | 1522.963525 | 1505.503889 | 1509.216744 | 1526.17316 | 1498.935264 |
| 19 | 1403.238282 | 1392.874059 | 1374.000767 | 1380.147497 | 1368.788918 | 1339.730679 |
| 20 | 1165.834444 | 1174.224599 | 1119.234628 | 1130.230438 | 1075.764944 | 1061.513324 |
| 21 | 1376.839049 | 1346.59496 | 1356.120547 | 1351.035138 | 1354.33679 | 1339.205461 |
| 22 | 1450.201446 | 1470.562769 | 1447.243032 | 1455.111487 | 1449.243032 | 1451.243032 |
| 23 | 1304.565098 | 1276.078273 | 1306.565098 | 1301.52575 | 1271.731385 | 1256.699807 |
| 24 | 1177.270794 | 1159.628041 | 1067.154873 | 1072.596415 | 1194.738271 | 1048.11836 |

| FEP | RW | RW PW | PH | PH PW | PH est-PW | PH est-PW2 |
| --- | --- | --- | --- | --- | --- | --- |
| 1 | 1337.336039 | 1323.424009 | 1333.477043 | 1329.795382 | 1309.051073 | 1297.184236 |
| 2 | 1436.078401 | 1439.21773 | 1427.911316 | 1431.904186 | 1426.615321 | 1422.032703 |
| 3 | 1590.77036 | 1588.709038 | 1628.808557 | 1626.40692 | 1629.934422 | 1631.762198 |
| 4 | 1646.589921 | 1649.704892 | 1728.042044 | 1729.412481 | 1751.465964 | 1721.001683 |
| 5 | 1666.149946 | 1666.149946 | 1762.81542 | 1762.81542 | 1764.81542 | 1766.81542 |
| 6 | 1428.691703 | 1431.354481 | 1430.691703 | 1428.07835 | 1421.24178 | 1427.073687 |
| 7 | 1288.751412 | 1291.092155 | 1288.083017 | 1293.160018 | 1283.907237 | 1284.560394 |
| 8 | 1363.419572 | 1377.857231 | 1358.128818 | 1364.643467 | 1360.128818 | 1361.966628 |
| 9 | 1646.366529 | 1645.064363 | 1727.644902 | 1724.733285 | 1729.644902 | 1735.91633 |
| 10 | 1148.701131 | 1153.331012 | 1023.516724 | 1042.152723 | 1189.812868 | 1016.488433 |
| 11 | 1089.374501 | 1074.669775 | 1029.702512 | 1034.386594 | 1016.300941 | 994.0407177 |
| 12 | NA | NA | NA | NA | NA | NA |
| 13 | 1520.830185 | 1529.489462 | 1526.07187 | 1528.26126 | 1514.987866 | 1530.07187 |
| 14 | 1706.240874 | 1696.333095 | 1777.905842 | 1796.521942 | 1768.241629 | 1775.819821 |
| 15 | 1375.44904 | 1351.505261 | 1329.875442 | 1326.38359 | 1323.585249 | 1271.012053 |
| 16 | 1463.618278 | 1484.279501 | 1465.077802 | 1467.696434 | 1468.800143 | 1469.077802 |
| 17 | 1404.06655 | 1399.365485 | 1398.828494 | 1397.450045 | 1423.002591 | 1410.014611 |
| 18 | 1307.087824 | 1317.75078 | 1286.92697 | 1295.222605 | 1311.319803 | 1290.66307 |
| 19 | 1457.817994 | 1455.173041 | 1455.755556 | 1456.65181 | 1456.665126 | 1458.138688 |
| 20 | 1654.010318 | 1639.070111 | 1741.23386 | 1736.864671 | 1751.49411 | 1741.058027 |
| 21 | 1438.753208 | 1478.64587 | 1439.849482 | 1441.309151 | 1441.849482 | 1444.753208 |
| 22 | 1439.898076 | 1432.719857 | 1421.165224 | 1421.000947 | 1456.411932 | 1411.40563 |
| 23 | 1486.446913 | 1496.16131 | 1488.446913 | 1487.487344 | 1490.446913 | 1483.688269 |
| 24 | 1597.856797 | 1596.023526 | 1641.405379 | 1633.401838 | 1637.888303 | 1635.524519 |
| 25 | 1270.786727 | 1277.20352 | 1241.998646 | 1238.874235 | 1291.807762 | 1247.186597 |
| 26 | 1760.854627 | 1768.761094 | 1931.179298 | 1967.931288 | 1921.039939 | 1893.122146 |

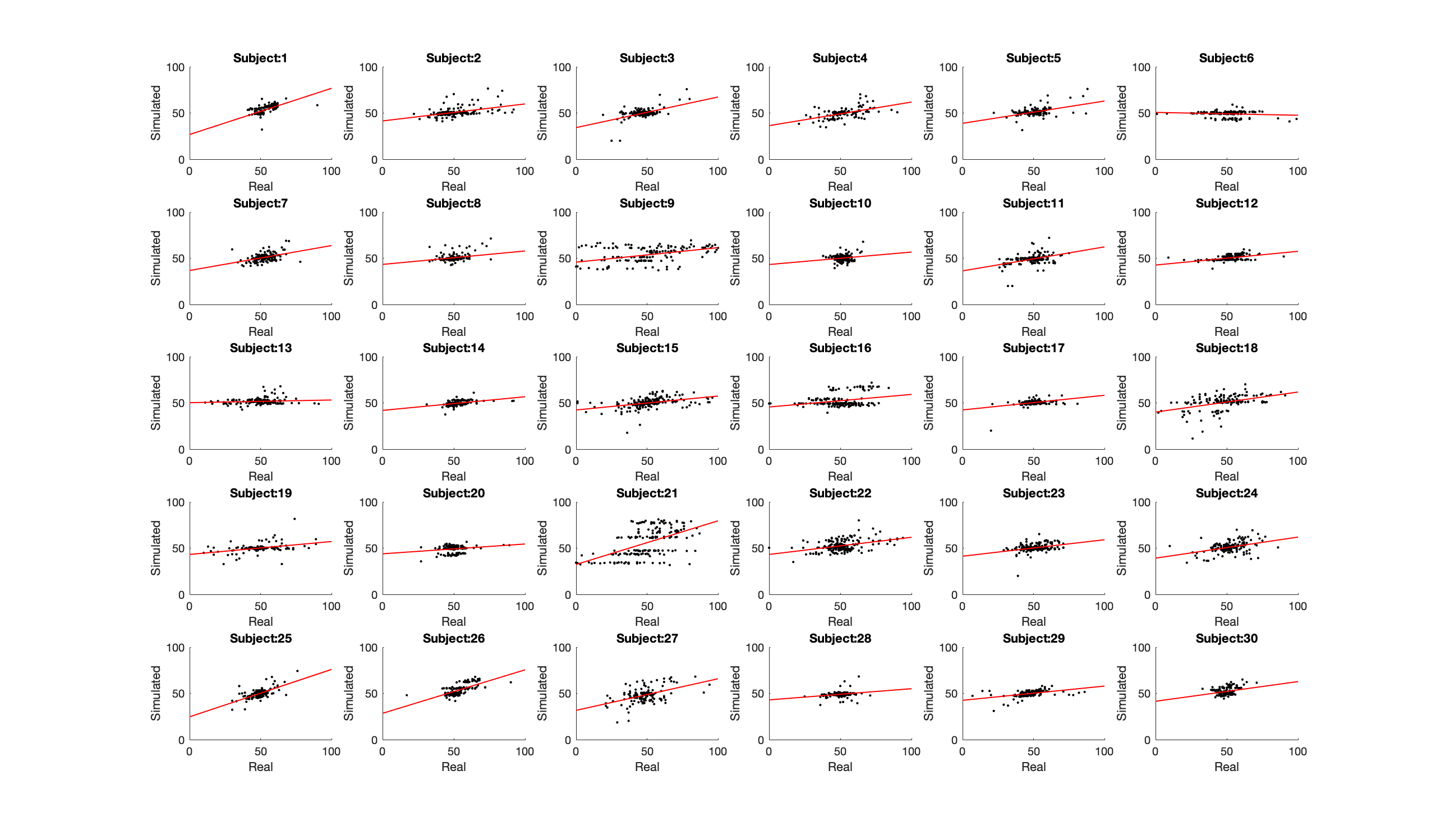

*Supplementary figure 4: Scatter plots, displaying the simulated data from the winning model in the healthy control group and the behavioural data.*

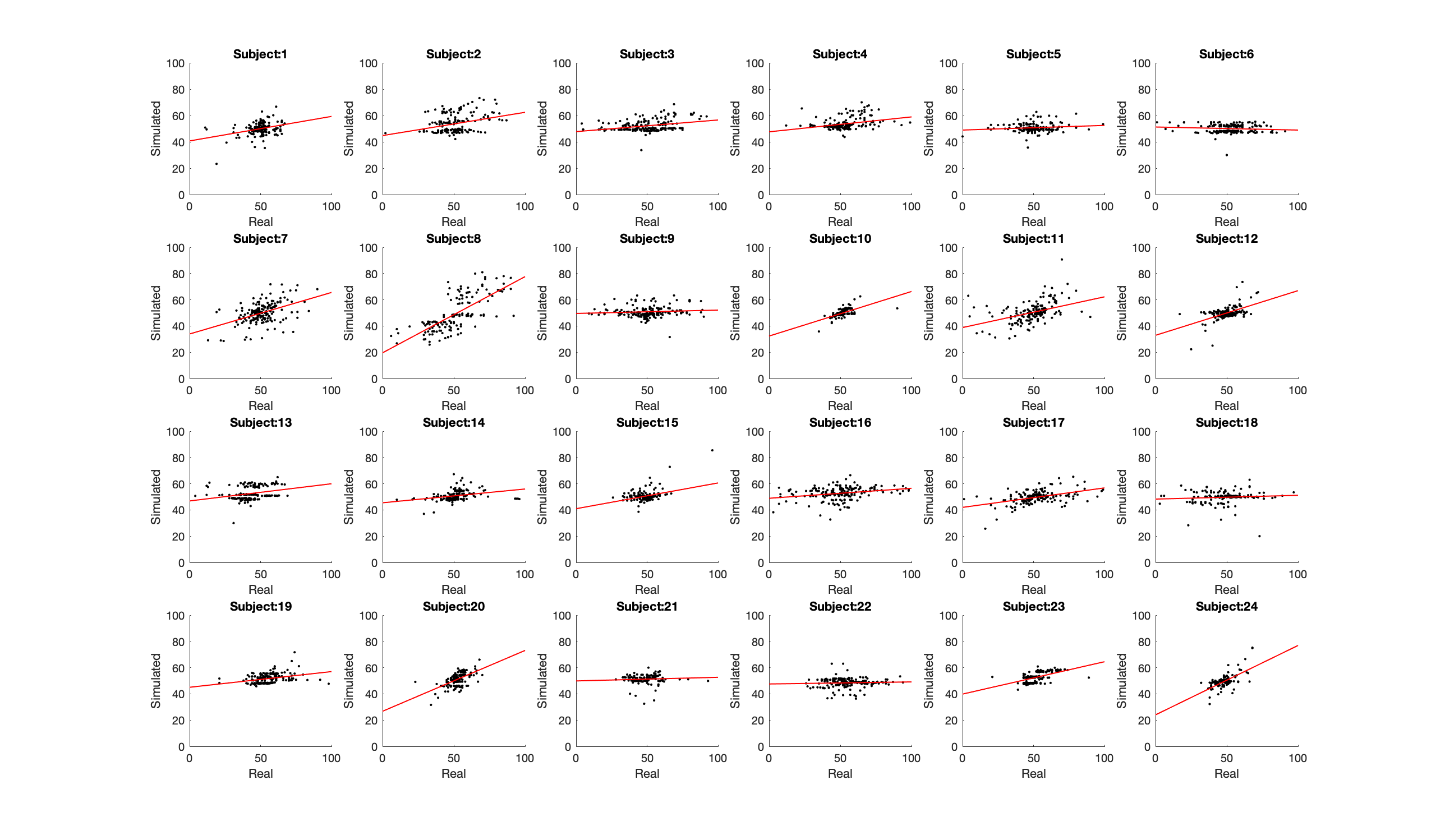

*Supplementary figure 5: Scatter plots, displaying the simulated data from the winning model in the ARMS group and the behavioural data.*

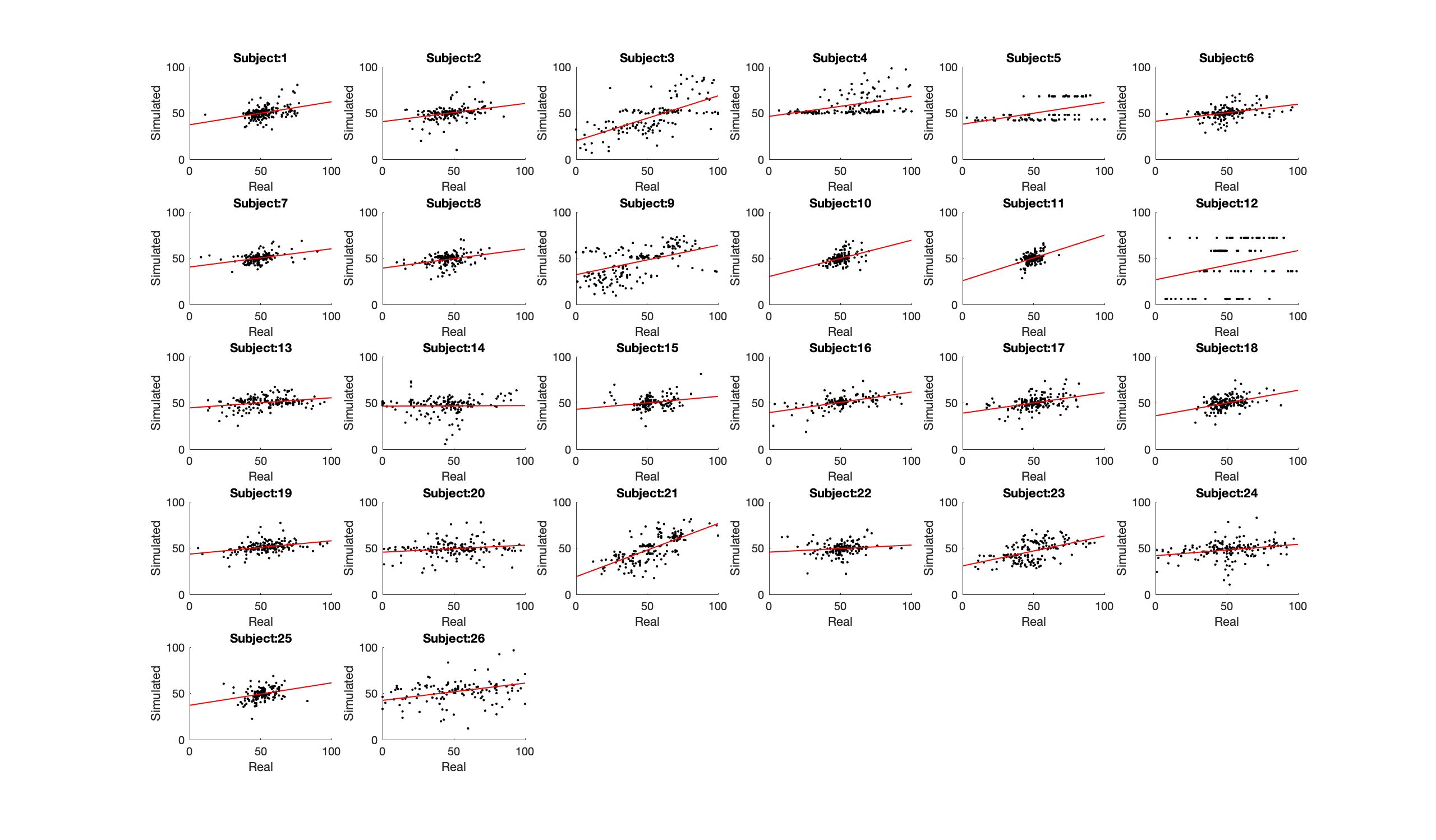

*Supplementary figure 6: Scatter plots, displaying the simulated data from the winning model in the FEP group and the behavioural data.*

*Elimination of possible confounding by effects on motivation*

In order to test whether there were possible confounding effects of motivation in the psychosis study we tested for differences in scrolling distance, missed trials and reaction times. We found a trend-level difference in missed trials (F{2,74}=2.93, *p*=.06), but not for scrolling distance and reaction times (all *p*>1).

*Comparison of model parameters between high and low precision conditions*

We found that both the HCS group T{29}=4.26, *p*<.001) and the ARMS group T{22}=4.99, *p*< .001) presented with significantly higher learning rates in the high precision condition compared to the low precision condition. By contrast, the FEP group did not show such a significant difference between high and low precision conditions (T{18}=1.5, *p*=.15). However, when we tested the precision by group interaction using a mixed (3x2) two-way ANOVA we did not see a significant interaction (F{2,76}=1.68 , *p*=.193) (Fig. 5G). We found no differences in learning rate decay between precision conditions for the HC group (T{29}=1.52, *p*= .14), ARMS group (T{22}=-.26, *p*= .74), and FEP group (T{18}=-1.39, *p*= .18) (Fig. 5H).

Supplementary Table 4: Psychosis study, significant clusters precision-weighting of unsigned prediction error

| Region | MNI | N-voxels | T | *p* |
| --- | --- | --- | --- | --- |
| Medial parietal lobe | [-9 -61 48] | 383 | 5.59 | <.001 |
| Occipital | [-12 – 101 2] | 87 | 5.29 | .011 |
| Right SFC | [24 12 58] | 171 | 4.53 | <.001 |
| Right PFC | [36 52 24] | 171 | 4.46 | <.001 |
| Left SFC | [-21 -2 52] | 94 | 4.20 | .007 |

*No evidence for an effect of antipsychotic medication on precision-weighting in patients*

We found no significant difference between FEP participants who take antipsychotic medication and those who do not in terms of precision weighting of the prediction error (T{18}=.14, p=.89; analysis conducted on voxels that showed a group difference). There was no significant relationship between anti-psychotic medication dosage and degree of precision-weighting in the 14 participants who took antipsychotic medications (r=.17, p=.52).

*Group differences in model fits do not drive results*

We added the r-values described above to the key behavioural and neural analyses, and show that all the effects remain significant. That is the group by precision condition interaction effect on behaviour remains significant when controlling for individual model fits as measured by the correlation between behavioural and simulated data (F{2,70}=3.73, *p*=.029). We extracted the betas from the small volume corrected ROI that showed differences between the FEP and healthy control group, and recomputed the group ANOVA both without model fit as a covariate (F{2,73}=5,65, *p*=.005) and with model fit as a covariate (F{2,73}=6.24, *p*=.003) and show that the effect strengthens after including model fit. Hence individual model fits do not appear to drive group differences on the behavioural and neural level. Furthermore, when we control for model fit in the analyses correlating symptoms with degree of precision-weighting, the effects remain the same (r=-.32, *p*=.023).

*Symptom correlations – additional analysis*

In order to explore the relationship between precision-weighting of prediction error and positive symptoms we performed an additional analyses where we included an additional 6 FEP patients (mean age: 26.4, 2 males, 4 females), who were not symptomatic, i.e. who did not have a score of >2 on P1 or P3 on the PANSS and so who did not meet eligibility for the main study r=0.-34, p=0.015, controlling for group, r=-0.28, p=0.054); within group FEP r=-0.35, p=0.08; within group ARMS r=-0.23, p=0.3. When controlling the correlation between symptoms and brain signal for model fit, the relationship is not changed: r=-0.32,p=0.02; IQ, r=-0.31, p=0.035 (1 participant had a missing IQ score). There is a marginal relationship between the degree of fMRI signal precision weighting and negative symptom severity (r=0.24, p=0.09). To further explore whether precision-weighting of prediction error is more correlated with specific types of psychotic symptoms we computed exploratory analyses where we correlated the degree of precision-weighting with different items of the PANSS, finding a correlation with P3 (hallucinatory behaviour) of r=-.34, p=.017, whereas other subscales did not correlate significantly, p>.05. However, we caution against over-interpretation of this as specific to hallucinations rather than delusions, as the P3 PANSS hallucinations item anchor points builds in delusional interpretation of the hallucination into scores of above 4 on this 7 point scale.

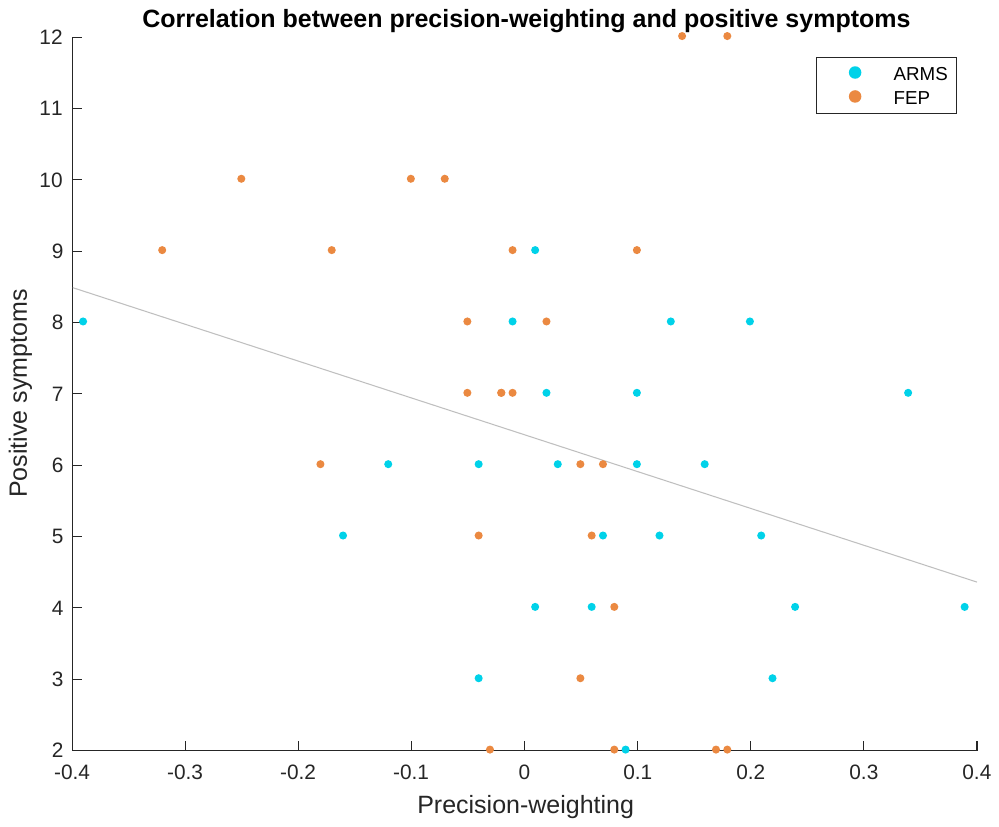

*Supplementary figure 7: Correlation between positive symptoms and precision-weighting for all patients, regardless of presence of symptoms.*

**Supplementary Discussion**

Our findings are in line with the wide literature on adaptive coding of neural signals to context. Adaptive coding seems to be a feature of many neural systems (Brenner et al., 2000; Fairhall et al., 2001; Barlow et al., 1961; Laughlin et al., 1981), ranging from the perceptual system (Ohzawa et al., 1982; Smirnakis et al., 1997) to reward systems (Tobler et al., 2005; Park et al., 2012). Indeed, previous studies that used the present paradigm have framed these questions in terms of adaptive coding of signed reward prediction errors rather than precision-weighting of unsigned prediction errors (Diederen et al., 2016 & 2017). However, adaptive coding of neural signals to the statistics of the environment follows directly from predictive coding models of neural functioning which suggest that prediction errors should be weighted by their precision (Srinivasan et al., 1982). Here the adaptive coding of prediction errors equals precision-weighting. Therefore, we believe that the framework of adaptive coding and precision-weighting do not constitute competing interpretations of the data, but rather that adaptive coding is a feature that follows from predictive coding models.

A limitation of the pharmacological study is that we only used one dose of each agent, which makes it hard to determine if the observed effects are dominated by presynaptic autoreceptors or postsynaptic receptors. It would be preferable to conduct a study with multiple assessments over the entire dose-range but this is not practical in human fMRI. Thus, whilst the pharmacological study indicates dopaminergic modulation of the processes under study, we cannot confidently infer that bromocriptine and sulpiride respectively represent enhanced and decreased dopaminergic signalling, and we do not wish to imply that either drug is a preclinical model of psychosis. We note that whilst sulpiride is a selective dopamine D2 receptor antagonist at the dose used, it does have actions on 5-HT1A at high doses. Bromocriptine is relatively selective for dopamine D2 receptors but also has action on some serotonergic receptors (Kvernmo et al., 2008).

Although we demonstrate that in health, dopaminergic modulation influences precision weighting of the prediction error signal in the superior cortex, and we demonstrate abnormal SFC precision weighting in psychosis, we cannot definitively conclude that the abnormalities in patients were of dopaminergic origin, as we make no neurochemical measures in patients. Such inference would require either a pharmacological design in patients, or the use of neurochemical imaging such as PET. One possible confound in the patient study is the use of anti-psychotic medication in some patients, given that we showed that precision-weighting of prediction errors is diminished after healthy controls take a dopamine D2-receptor antagonist in controls. We addressed this issue by showing that there are no differences in precision-weighting within the participants who take anti-psychotic medicaitons and those who do not. We furthermore showed that there is no relationship between anti-psychotic dosage and the degree of precision-weighting in this study. Furthermore we demonstrated evidence for aberrancies in cortical precision-weighting of prediction errors, and impaired task performance, relating to schizotypal personality in the healthy population, where medication cannot be a confound. We thus conclude that there is no evidence in the present study that the reported effects in patients are driven by the use of anti-psychotic medications.

In the present study, no differences were found with regards to the ARMS group. This is in line with recent work showing intact cortical prediction error responses in ARMS (Ermakova et al., 2018). There might be different reasons for this. First, considering that the effects were correlated with severity of symptoms, and the symptoms in the ARMS group were significantly lower, the lack of a significant finding for the ARMS group might be a matter of severity. Indeed, if we separate the ARMS group and FEP group and correlate the degree of precision-weighting with positive symptoms, we find comparable r values for both groups (ARMS: r=-.30, FEP: r=-.36), although neither is significant in isolation due to the small N. An alternative is that the aberrancies are disease category specific, which could be relevant as only a minority of ARMS patients will go on to develop psychotic illness (Morrison et al 2012 BMJ). Future studies with more power, ideally including longitudinal follow-up, would clarify these issues.

**References specific to supplementary material**

Ashburner, J., & Friston, K. J. (2005). Unified segmentation. *Neuroimage*, *26*(3), 839-851.

Barlow, H. B. (1961). Possible principles underlying the transformations of sensory messages.

Brenner, N., Bialek, W., & Van Steveninck, R. D. R. (2000). Adaptive rescaling maximizes information transmission. *Neuron*, *26*(3), 695-702.

Crespi LP: Quantatative variation of incentive and performance in the white rat. *American Journal of Psychology*. 1942, 55: 467-517. 10.2307/1417120.

Fairhall, A. L., Lewen, G. D., Bialek, W., & van Steveninck, R. R. D. R. (2001). Efficiency and ambiguity in an adaptive neural code. *Nature*, *412*(6849), 787.

Kvernmo, T., Houben, J., & Sylte, I. (2008). Receptor-binding and pharmacokinetic properties of dopaminergic agonists. *Current topics in medicinal chemistry*, *8*(12), 1049-1067.

Laughlin, S. (1981). A simple coding procedure enhances a neuron's information capacity. *Zeitschrift für Naturforschung c*, *36*(9-10), 910-912.

Morrison AP, French P, Stewart SL, Birchwood M, Fowler D, Gumley AI, Jones PB, Bentall RP, Lewis SW, Murray GK, Patterson P, Brunet K, Conroy J, Parker S, Reilly T, Byrne R, Davies LM, Dunn G (2012), “Early detection and intervention evaluation for people at risk of psychosis: multisite randomised controlled trial.” *BMJ* 344:e2233

Niv Y: Cost, benefit, tonic, phasic: what do response rates tell us about dopamine and motivation? *Ann N Y Acad Sci.* 2007, 1104: 357-376. 10.1196/annals.1390.018.

Ohzawa, I., Sclar, G., & Freeman, R. D. (1982). Contrast gain control in the cat visual cortex. *Nature*, *298*(5871), 266.

Park, S. Q., Kahnt, T., Talmi, D., Rieskamp, J., Dolan, R. J., & Heekeren, H. R. (2012). Adaptive coding of reward prediction errors is gated by striatal coupling. *Proceedings of the National Academy of Sciences*, 201119969.

Poser, B. a., Versluis, M. J., Hoogduin, J. M., & Norris, D. G. (2006). BOLD contrast sensitivity enhancement and artifact reduction with multiecho EPI: Parallel-acquired inhomogeneity-desensitized fMRI. *Magnetic Resonance in Medicine*, *55*(6), 1227–1235.

Raine, A. (1991). The SPQ: a scale for the assessment of schizotypal personality based on DSM-III-R criteria. *Schizophrenia bulletin*, *17*(4), 555.

Smirnakis, S. M., Berry, M. J., Warland, D. K., Bialek, W., & Meister, M. (1997). Adaptation of retinal processing to image contrast and spatial scale. *Nature*, *386*(6620), 69.

Srinivasan, M. V., Laughlin, S. B., & Dubs, A. (1982). Predictive coding: a fresh view of inhibition in the retina. *Proc. R. Soc. Lond. B*, *216*(1205), 427-459.

Tobler, P. N., Fiorillo, C. D., & Schultz, W. (2005). Adaptive coding of reward value by dopamine neurons. *Science*, *307*(5715), 1642-1645.
